# Metabolic control of smooth muscle cell phenotype switching in atherosclerosis

**DOI:** 10.64898/2026.05.19.726223

**Authors:** Rong-Mo Zhang, Xiaolong Zhu, Hosung Bae, Jiasheng Zhang, Yanming Li, Pei-Yu Chen, Ying H. Shen, George Tellides, Nathaniel Snyder, Cholsoon Jang, Martin A. Schwartz, Zoltan Arany, Michael Simons

## Abstract

The loss of smooth muscle cell (SMC) contractile phenotype contributes to various diseases including atherosclerosis. However, its metabolic basis is not entirely elucidated. Since the transforming growth factor beta (TGFβ) signaling is among principal regulators of SMC contractility, we studied metabolic regulation of TGFβ signaling in SMCs in vitro and atherosclerotic mouse models and human lesions. We found that TGFβ induced Ac-CoA synthetase 2 (ACSS2)-dependent Ac-CoA production, by suppressing pyruvate dehydrogenase kinase 4 (PDK4). This stabilized R-SMADs and TGFβ receptor 1, preserving SMC contractile phenotype. SMC-specific PDK4 knockout mimicked the effect of TGFβ signaling both metabolically and phenotypically, increasing glucose-derived synthesis of Ac-CoA and SMC contractile phenotype. SMC-specific *Pdk4* knockout in *ApoE* knockout mice reduced atherosclerosis. Furthermore, human specimens demonstrated a strong correlation between PDK4 level and atherosclerosis severity. These findings indicate that continuous TGFβ signaling, critical to the maintenance of the normal SMC contractile state and is regulated by PDK4 and carbohydrate metabolism.

**Teaser:** Reducing PDK4 metabolically restricts aortic plaque growth via TGFβ-dependent SMC contractility.

## INTRODUCTION

Atherosclerosis is a complex disease characterized by development of intravascular atherosclerotic plaques that gradually compromise the arterial lumen and become prone to rupture (*1, 2*). A number of cell types, including smooth muscle cells (SMCs), endothelial cells and various inflammatory cells contribute to plaque development (*3, 4*). Key changes include the process of endothelial-to-mesenchymal transition (EndMT), a series of events that result in the loss of endothelial cell identity of the endothelium lining the plaque surface and the expansion of the mass of SMC-like cells (*5*). In parallel, native SMCs lose their differentiated, contractile state, switching to a proliferative phenotype thereby contributing to the mass of cells in the developing plaque (*4, 6*). While the SMC phenotype switch is well described, molecular events controlling this process have not been fully established. Given the long course of this SMC cell fate transition, we hypothesized that a metabolic switch is the underlying molecular driver. Since TGFβ is thought to play a key role in the maintenance of SMC contractile state (*7*) and given our recent description of the metabolic control of TGFβ signaling in the endothelium (*8*), we set out to study a potential metabolic switch underlying SMC phenotype switch in atherosclerosis settings.

Transforming growth factor β (TGFβ) is a pleiotropic growth factor with various complex effects in different cells and tissues. Dysregulation of TGFβ signaling is associated with multiple diseases, including atherosclerosis (*7, 9–11*), aneurysms (*12–15*), and cancer (*16–18*) among others. Importantly, while many effects of TGFβ signaling are cell-type specific, the molecular basis of this specificity has not been established. In the vasculature, TGFβ signaling is critical to normal smooth muscle cell (SMC) homeostasis while it destabilizes endothelial cells (ECs) (*6, 19, 20*). Previous studies from our laboratory have identified a cell type-specific role of TGFβ signaling in atherosclerosis with TGFβ having a pro-inflammatory, pro-atherosclerotic effect in ECs (*10*) and anti-inflammatory, anti-atherosclerotic effects in SMCs (*9*).

TGFβ signaling involves both canonical and non-canonical pathways. The canonical signaling involves SMAD adaptor proteins while the non-canonical signaling utilizes various intracellular kinases (*6, 12, 21*). A particular interesting feature of canonical TGFβ signaling is its slow onset of action with up to 7 days required to observe phenotypic changes in both ECs and SMCs (*9, 10*). This long duration of action suggests a potential involvement of metabolic reprogramming. This has been described in endothelial cells where TGFβ induces activation of glycolysis and increases production of glucose-derived Acetyl-CoA (Ac-CoA) that results in acetylation and stabilization of key canonical TGFβ signaling molecules and enhancement of TGFβ signaling (*8*). The critical step was TGFβ-dependent inhibition of expression of pyruvate dehydrogenase kinase 4 (PDK4) that allowed for PDH-dependent synthesis of acetate(*22*) followed by Ac-CoA synthetase 2 (ACSS2)-mediated Ac-CoA synthesis in the cytoplasm. The end effect was activation and dedifferentiation of ECs, a process referred to as endothelial-to-mesenchymal transition (EndMT).

In contrast to dedifferentiating effects of TGFβ signaling in EC (*8, 10*), the same canonical TGFβ signaling induces a differentiated, contractile phenotype in SMCs (*7, 9*). Since the metabolic aspects of TGFβ signaling in SMCs have not been sufficiently explored, we set out to study whether metabolic reprogramming is involved in regulation of TGFβ signaling in SMCs and whether a difference in metabolic controls can explain cell-type (EC vs. SMC) specific effects of TGFβ signaling. Surprisingly, we observed that TGFβ signaling in SMC induced very similar metabolic perturbations to that observed in EC. In particular, there was an increase in SMC glucose uptake due to increased expression of the glucose transporter GLUT1, decreased PDK4 expression and activation of PDH/ACSS2-dependent Ac-CoA synthesis in the cytoplasm. The result was an increase in SMC contractile state. To assess the biological significance of these observations, we induced an SMC-specific deletion of *Pdk4* in mice on an atherosclerotic *ApoE* null background challenged by either high fat diet or complete carotid ligation. In agreement with the *in vitro* and metabolic data, this resulted in increased maintenance of the SMC contractile phenotype and a significant reduction in the development and progression of atherosclerosis. These findings indicate a complex nature of metabolic control of TGFβ signaling and demonstrate that the same metabolic chain of events may lead to very different outcomes in endothelial cells and SMCs cell types.

## RESULTS

### TGFβ increases glycolysis and contractility of smooth muscle cells

To evaluate the effects of TGFβ signaling in SMCs, we carried out RNAseq analysis of primary human aortic SMC (HASMCs) after 7 days of TGFβ stimulation (Fig. 1A). As expected, most contractile genes, including *ACTA2*, *CALD*, *CNN1*, *MYL9* and *TAGLN*, were upregulated, while SMC dedifferentiation marker *KLF4* was decreased (Fig. 1A and supplementary Fig. 1A). Western blotting confirmed activation of canonical TGFβ signaling (SMAD2/3 phosphorylation, Supplementary Fig 1B) while immunocytochemistry showed translocation of SMAD2/3 into the nucleus as well as increased SMA staining and decreased proliferation as measured by the Ki67 marker expression (supplementary Fig. 1C). There were also pronounced changes in expression of genes associated with cellular metabolism. In particular, expression of genes promoting glycolysis such as *SLC2A1*, *PFKFB3*, *HK1* was increased (Fig. 1A).

**Figure 1.**
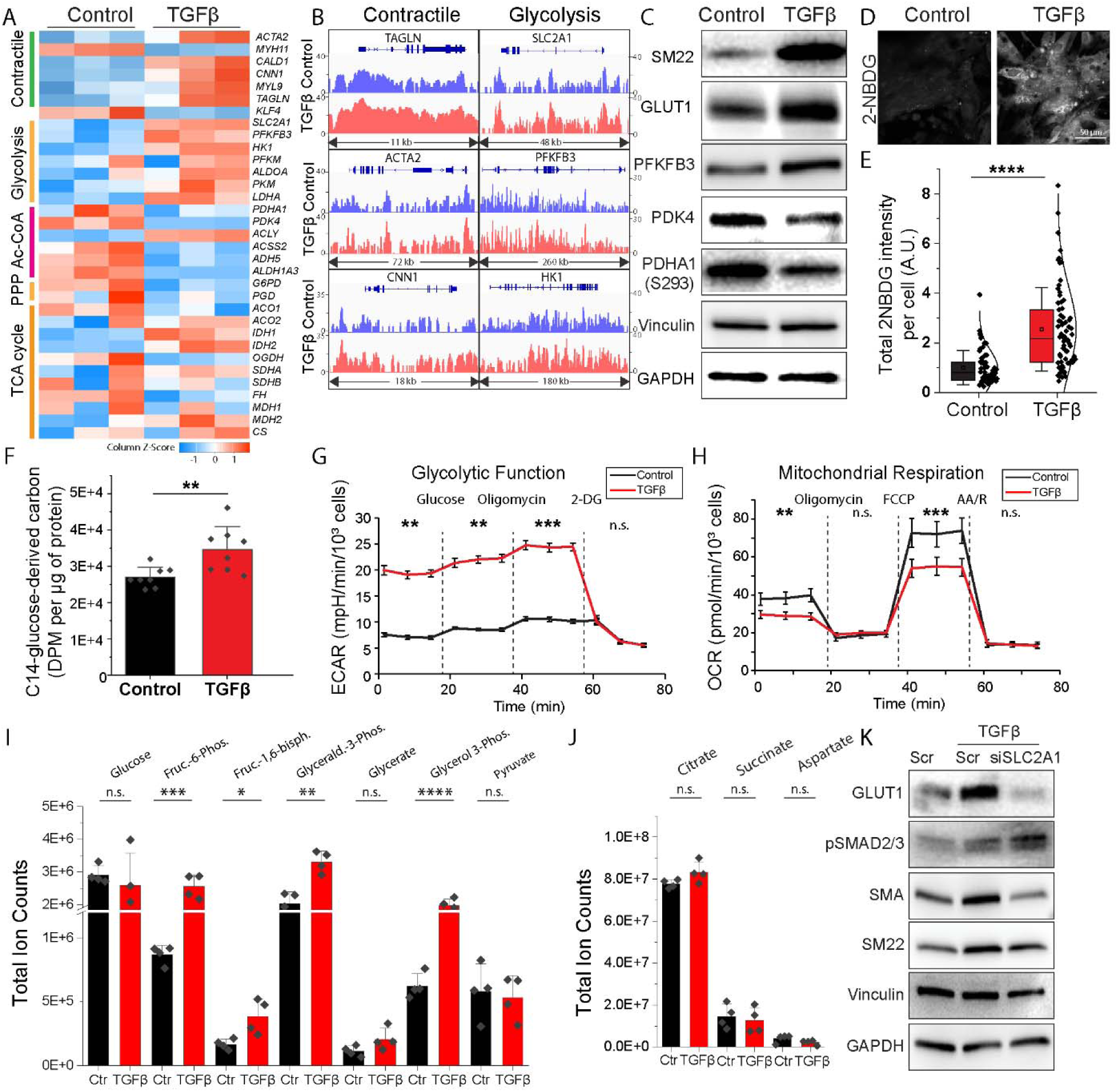
Metabolic effects of HASMC TGF-β signaling. (**A**) Bulk RNA-seq analysis of SMC contractile and metabolic gene expressions of glucose oxidation after 7d TGFβ treatment. Note that expression of genes involved in glycolysis is increased. (**B**) ATAC-seq of key genes involved in HASMC contractile phenotype and glycolysis after 7d TGFβ treatment. (**C**) Protein levels of SMC contractile and metabolic gene after TGFβ treatment. (**D**) Glucose uptake assay using the fluorescent tracer 2-(N-(7-Nitrobenz-2-oxa-1,3-diazol-4-yl) Amino)-2-Deoxyglucose (2-NBDG). 2NBDG was added for 3 hours after 7 days of TGFβ treatment of HASMCs. (**E**) Quantification of cellular 2-NBDG intensity. Each dot represents one cell. (**F**) Quantification of C^14^ radioactivity in CO_2_ produced by HASMCs fed with C^14^-glucose (D-[14C(U)]-glucose) for 24 h after 7 days of TGFβ stimulation. Disintegrations per minute (DPM) of C^14^ radioactivity was normalized to 1 µg of protein of cell lysate. (**G-H**) Seahorse analysis of glycolytic function (ECAR, g) and mitochondrial respiration (OCR, h), after 7 days of TGFβ treatment. Note that TGFβ treatment increased ECAR and decreased OCR of HASMCs. One-way ANOVA was used to compare treatment at each metabolic phase. (**I-J**) Liquid chromatography-mass spectrometry (LC-MS) metabolomics analysis of key metabolites of glycolysis (i) and the TCA cycle (j) in HASMCs after 7 days of TGF-β stimulation. Total ion counts were normalized to 10^6^ cells. (**K**) Western blotting analyzing the effects of GLUT1 KD (siSLC2A1) on HASMC contractile phenotype (αSMA and SM22) under 10 ng/mL TGFβ treatments for 3 days.

In agreement with the above, ATAC-seq analysis demonstrated open conformation of the key SMC contractile and glycolytic genes (Fig. 1B) following TGFβ treatment. Western blotting confirmed increased expression of SMC contractile gene marker SM22, the glucose transporter GLUT1 (*SLC2A1*) and the glycolytic regulator PFKFB3 by TGFβ (Fig. 1C). Consistent with the increase in GLUT1 expression, fluorescent tracer assay using 2-(N-(7-Nitrobenz-2-oxa-1,3-diazol-4-yl) Amino)-2-Deoxyglucose (2NBDG) showed increased glucose uptake (2.8-fold) by HASMCs exposed to TGFβ (Fig. 1D&E). Decreased pyruvate dehydrogenase lipoamide kinase isozyme 4 (PDK4) and phosphorylation of pyruvate dehydrogenase α (PDHA1, S293) are also consistent with elevated glucose metabolism (Fig. 1C and supplementary Fig. 1D) as is increased CO_2_ generation from the C^14^-labelled glucose (Fig. 1F).

Seahorse analysis showed an increase in the extracellular acidification rate (ECAR) after 7 days of TGFβ treatment (Fig. 1G and supplementary Fig. 1E) while the oxygen consumption rate (OCR) decreased (Fig. 1H and supplementary Fig. 1F). These results agree with the RNAseq data, indicating that TGFβ exposure switches SMC metabolism into a more glycolysis-dependent energy production mold. Detailed mass spectrum measurements of metabolites involved in glycolysis and TCA cycle showed substantial increases in the total fructose-6-phosphate, fructose-1,6-biphosphate, glyceraldehyde-3-phosphate and glycerol 3-phosphate levels (2.9, 2.3, 1.6 and 3.2-fold respectively, Fig. 1I). Furthermore, there was an increase in ^13^C-glucose-derived enrichment of glycolytic intermediates after TGFβ treatment (supplementary Fig. 1G-J), while TCA cycle intermediates were not altered in total amount and tended to be less ^13^C-enriched (Fig. 1J and supplementary Fig. 1H).

To study whether TGFβ-increased glycolysis contributes to the induction of the SMC contractile phenotype, we knocked down GLUT1, one of the most abundant glucose transporters in HASMCs. This resulted in a reduction in contractile genes expression in the presence of TGFβ stimulation (and supplementary Figs. J-K). In the absence of TGFβ stimulation or in the presence of the ALK5 inhibitor, GLUT1 KD further reduced expression of contractile SMC genes (Fig. 1K and supplementary Figs. 1K-R). These results indicate that TGFβ stimulation increases SMC glucose consumption and that both TGFβ signaling and glucose consumption play a role in regulation of SMC contractile gene expression.

### TGFβ-induced suppression of PDK4 expression mediates SMC contractile protein expression and glycolysis

To further clarify metabolic effects of canonical TGFβ signaling in HASMCs, we focused on the pyruvate dehydrogenase kinase 4 (PDK4), a direct SMAD2/3 target (*8*). Analysis of the Human Protein Atlas(*23*) showed that among the four PDK isozymes, PDK4 is by far the most abundant isoform in human aortic SMCs in vivo (supplementary Fig. 2A). This was confirmed by qPCR analysis of primary human aortic SMCs in culture (Fig. 2A). In line with previous results in endothelial cells (ECs) (*8*), *PDK4* expression was reduced by 95% following SMC treatment with TGFβ (Fig. 2A).

**Figure 2.**
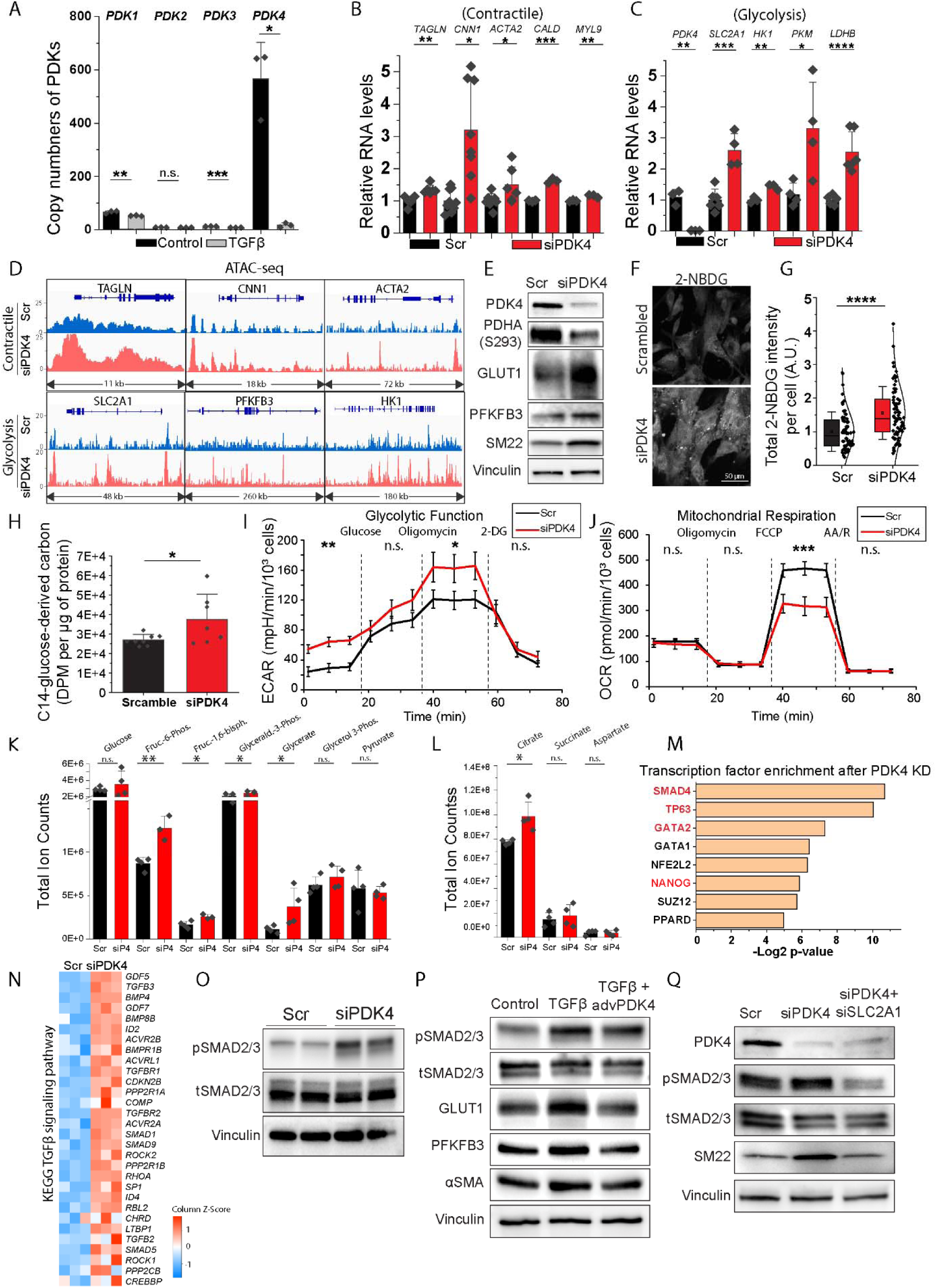
Metabolic effects of PDK4 knockdown in HASMCs. (**A**) Copy number of all four PDK4 isoforms (*PDK1*, *PDK2*, *PDK3* and *PDK4*) from bulk RNA-seq of HASMCs after TGFβ treatment for 7 days. (**B-C**) qPCR analysis of key markers for HASMC contractile phenotype (B) and glycolysis (C), after PDK4 KD treatment for 7 days. (**D**) ATAC-seq analysis of key genes involved in HASMC contractile phenotype and glycolysis after 7d PDK4 KD treatment. (**E**) Western blotting of key enzymes involved in glycolysis (GLUT1 and PFKFB3) and HASMC contractile phenotype (SM22) after 7d PDK4 KD treatment. (**F-G**) 2-NBDG fluorescence and quantification of intensity after 3 hours incubation with HASMCs of PDK4 KD for 7 days. (**H**) Quantification of C^14^ radioactivity in CO_2_ produced by HASMCs fed with C^14^-glucose (D-[14C(U)]-glucose) for 24 h after 7 days of PDK4 KD treatment. DPM of C^14^ radioactivity was normalized to 1 µg of cell lysate protein. (**I-J**) Seahorse assays measuring glycolytic function, via ECAR and mitochondrial respiration, via OCR, after 7 days of PDK4 KD treatment. One-way ANOVA was used to compare treatment at each metabolic phase. (**K-L**) LC-MS metabolomics analysis of key metabolites in glycolysis (k) and the TCA cycle (l) HASMCs after 7 days of PDK4 knockdown treatment. Total ion counts were normalized to 10^6^ cells. (**M**) Transcription factor enrichment after PDK4 KD using ChIP Enrichment Analysis (ChEA). (**N**) Heatmap of top 30 most unregulated mRNAs in TGFβ signaling. (**O**) Western blotting of SMAD2/3 phosphorylation after PDK4 KD for 7 days. (**P**) Protein levels of SMAD2/3 phosphorylation, key glycolysis enzymes (GLUT1 and PFKFB3) and SMC contractile marker (αSMA) after 10ng/mL of TGFβ with PDK4 adenovirus or control virus for 3 days. (**Q**) Western blotting of SMAD2/3 phosphorylation and SMC contractile marker (SM22) after 7 days of PDK4 KD or PDK4 and GLUT1 double KD.

To investigate whether a PDK4 knockdown (KD) would mimic the effect of TGFβ treatment on SMC contractile gene expression and glycolysis, primary HASMCs were treated with the PDK4 siRNA (supplementary Fig. 2B). qPCR analysis of gene expression 7 days later showed increased expression of a number of contractile proteins including *TAGLN*, *CNN1*, *ACTA2*, *CALD* and *MYL9* (Fig. 2B) and glycolytic genes (Fig 2C). Concordantly, ATAC-seq revealed that the transposase accessibility to *TAGLN*, *CNN1*, *ACTA2*, as well as SLC2A1, PFKFB3 and HK1 was increased (Fig. 2D). At the protein levels, expression of SM22 was increased 7 days following PDK4 knockdown (Fig. 2E and supplementary Fig. 2C). Finally, immunostaining showed increased αSMA expression and decreased cellular proliferative activity measured by Ki67 expression (supplementary Fig. 2D), without changes in cell viability (supplementary Fig. 2E-F). Chemical inhibitors of PDKs, including DCA and PDK4-IN-1 (an anthraquinone derivative PDK4 inhibitor) had the same effect on SMAD2/3 phosphorylation and SM22 (Fig 2E and supplementary Fig. 2G-H). Western blots also confirmed increased expression of GLUT1 and PFKFB3 proteins (Fig. 2E) while glucose uptake was increased 1.6-fold, as indicated by the 2-NBDG assay (Fig. 2F-G).

Glucose consumption was also increased as shown by C^14^ glucose carbon tracing to ^14^CO_2_ following PDK4 knockdown (Fig. 2H). Concordantly, Seahorse assays showed that siPDK4 strongly induced ECAR and reduced OCR (Fig. 2I-J), with elevated glycolytic capacity reserve (supplementary Fig. 2I-J). Mass spectroscopy analysis of metabolites involved in glycolysis and the TCA cycle indicated that PDK4 KD increased total levels of key glycolytic metabolites including fructose-6-phosphate, fructose-1,6-biphosphate and glycerate (Fig. 2K). To better characterize the source of these metabolites, HASMCs were treated with C^13^-glucose 7 days after PDK4 KD. Mass spectrometry analyses indicated that the proportion of C^13^-containing metabolites, including fructose-6-phosphate, fructose-1,6-biphosphate, and pyruvate was increased by PDK4 KD (supplementary Fig. 2K) while most key metabolites in the TCA cycle, including succinate and aspartate, were not changed by PDK4 KD (Fig. 2L, supplementary Fig. 2L). Taken together, these data indicate that in SMCs the effects of PDK4 KD are similar in terms of induced metabolic changes and contractile state to TGFβ stimulation. Both interventions increased glucose-derived CO_2_ production while glucose utilization in the TCA cycle remained largely unchanged, suggesting an alternative usage of glucose carbon other then fueling the TCA cycle. In agreement with this possibility, carbon-tracing studies showed an increase in the pentose phosphate pathway activity after both TGFβ stimulation and PDK4 KD as shown by an increased percentage of glucose-derived C^13^ carbon in both ribose-5P and sedoheptulose-7P (Supplementary Figs. 1I-J and 2N-O).

To further elucidate the role of PDK4 in regulation of the SMC contractile phenotype, we examined changes in SMC gene expression 7 days following the PDK4 KD. Bulk RNAseq revealed 525 differentially expressed genes (fold change > 2, p < 0.05) following the PDK4 KD (supplementary Fig. 2M). ChIP Enrichment Analysis (ChEA) identified several transcription factors capable of regulating expression of differentially expressed genes identified in bulk RNAseq (supplementary Fig. 2M). Among these were transcription factors known to be involved in canonical TGFβ signaling including SMAD4, TP63 (*24*), GATA2 (*25*), and NANOG (*26*) among others (Fig. 2M). A heat map (Fig 2N) shows the top 30 upregulated genes involved in TGFβ signaling upon PDK4 knockdown with most of these involved in the TGFβ signaling cascade. Western blotting of HASMCs confirmed increased SMAD2/3 phosphorylation levels following PDK4 KD (Fig. 2O and supplementary Fig. 2S). Conversely, overexpression of PDK4 abrogated SMAD2/3 phosphorylation and SMC contractile gene expression following treatment with TGFβ (Fig. 2P and supplementary Fig. 2T-W). These data indicate that a reduction in PDK4 level serves as positive feedback for TGFβ signaling. Finally, SLC2A1 knockdown abrogated the increase in pSMAD2/3 following PDK4 KD (Fig. 2Q and supplementary Fig. 2X-Y), thus demonstrating that glucose utilization is necessary for PDK4 KD-dependent induction of TGFβ signaling in HASMCs.

### A reduction in PDK4 expression induces ALK5 and SMAD2/3 acetylation via increased Ac-CoA production

Mass spectroscopy measurements of total cellular Ac-CoA levels in primary HASMC showed a significant increase following TGFβ stimulation or PDK4 KD (Fig. 3A). Metabolic tracing using C^13^-glucose showed increased incorporation of the glucose-derived carbons into Ac-CoA and acetate were of the same order of magnitude as the observed by TGFβ stimulation or PDK4 knockdown (supplementary Fig. 2O-Q). We have previously reported that TGFβ signaling induces an increase in ALK5, SMAD2 and SMAD4 acetylation in endothelial cells (*8*). To see if this increase in Ac-CoA levels also affected acetylation of these key proteins in the SMC TGFβ signaling cascade, we performed immunoprecipitation of acetylated proteins upon either TGFβ stimulation or PDK4 KD. As in endothelial cells, acetylation ALK5 and SMAD2/3 (acetylated to total protein ratio) were increased both by TGFβ treatment and PDK4 KD (Figs. 3B-D and 3E-G).

**Figure 3.**
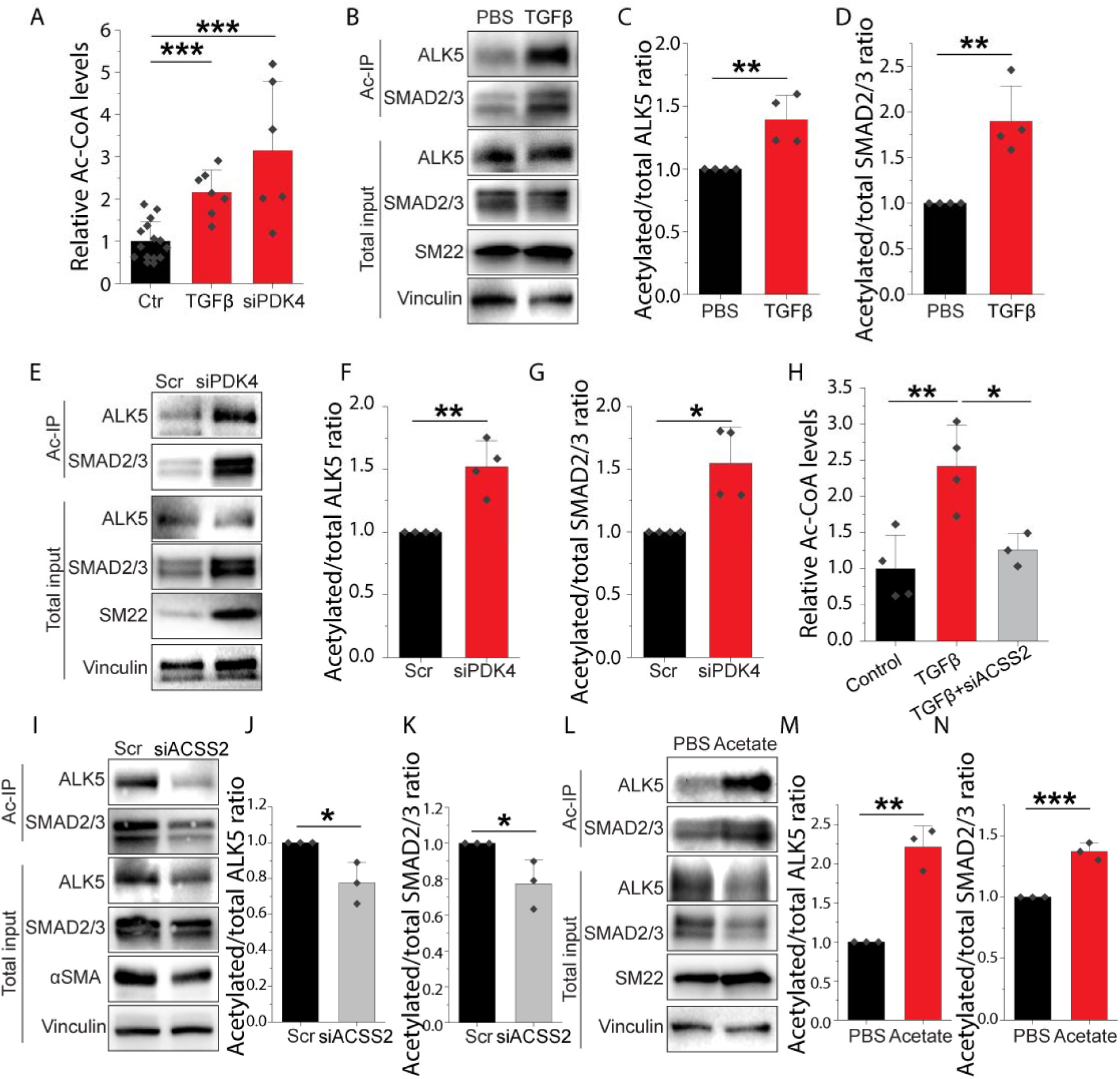
TGFβ and PDK4 knockdown induce ALK5 and SMAD2/3 acetylation via ACSS2-assisted Ac-CoA production. (**A**) Relative levels of Ac-CoA in HASMCs treated with TGFβ or PDK4 KD for 7 days. Ac-CoA levels were measured by liquid chromatography-high resolution mass spectrometry. (**B-G**) Immunoprecipitation of acetylated ALK5 and SMAD2/3 after 7 days of TGFβ treatment (B) or PDK4 KD (E). SM22 was used to present phenotype changes of HASMCs. Vinculin was used as loading control. C-D and F-G represent quantification of three-four independent experiments. (**H**) Relative levels of Ac-CoA in HASMCs treated with TGFβ or TGFβ + ACSS2 KD for 7 days. Note that ACSS2 knockdown brings back the elevated Ac-CoA levels by TGFβ treatment. (**I-N**) Immunoprecipitation of acetylated ALK5 and SMAD2/3 after 7 days of ACSS2 knockdown (I) or exogenous acetate treatment (E). αSMA or SM22 was used to present phenotype changes of HASMCs. Vinculin was used as loading control. Paired-sample t-tests were used to test statistical significance for (C, D, F, G, J, K, M and N). Note that the quantifications reflect the acetylated to total protein ratio. One-way ANOVA was used to assess statistical significance for (A&H).

Ac-CoA can be generated via the citrate shuttle by the enzyme ACLY and by a direct conversion of acetate by the enzyme ACSS2. The latter has been shown to play an important role in Ac-CoA generation in endothelial cells (*8*). Western blot analysis of the presence of acetylated lysine (Ac-K) showed that both TGFβ and PDK4 KD increased protein acetylation levels in both mitochondria and cytoplasm (supplementary Fig. 3A). A knockdown of either ACSS2 or ACLY reduced TGFβ signaling and SMC contractile gene expression (supplementary Fig. 3B). However, adenoviral overexpression of ACLY could not rescue the TGFβ signaling or SMC contractile gene expression reduced by ACSS2 KD (supplementary Fig. 3C). Together, these results indicate that TGFβ or PDK4 KD could increase protein acetylation in both mitochondria and the cytoplasm. However, ACSS2-derived cytoplasmic Ac-CoA plays a major role in TGFβ-induced SMC contractile gene expressions. To further validate ACSS2 involvement in SMC Ac-CoA generation, we carried out mass spectroscopy analysis of Ac-CoA levels in HASMC. A knockdown of ACSS2 abrogated TGFβ-induced increase in Ac-CoA level (Fig. 3H). The extent of ALK5 and SMAD2/3 acetylation was also reduced following ACSS2 KD (Figs. 3I-K). To further investigate the role of acetate in SMC TGFβ signaling cascade, we treated HASMCs with acetate for 48h and observed increased acetylation of ALK5 and SMAD2/3 (Figs. 3L-N).

### ACSS2 is essential for TGF**β** signaling in human aortic smooth muscle cells

To further analyze the role of ACSS2 in SMC TGFβ signaling, transcript levels of key genes involved in glucose metabolism were analyzed using RNAseq in the presence or absence of ACSS2 KD (Fig. 4A). Gene ontology analysis showed that glycolysis & gluconeogenesis pathways were the most downregulated KEGG pathways when ACSS2 was knockdown (Fig. 4B), with downregulation of several key regulators including *PFKFB3*, *PKM* and *LDHA* (Fig. 4C). Seahorse measurements showed that ACSS2 KD reduced the TGFβ-induced increase in glycolysis and reduction in OCR (Figs. 4D-E). Furthermore, ACSS2 KD prevented the elevation of glucose-derived CO_2_ production by TGFβ (Fig. 4F). Finally, similarly to TGFβ, acetate treatment of HASMC in culture increased glycolysis (Fig. 4G) while also increasing OCR (Fig. 4H). Acetate treatment also increased glucose-derived C^14^ incorporation into CO_2_ (Fig. 4I).

**Figure 4.**
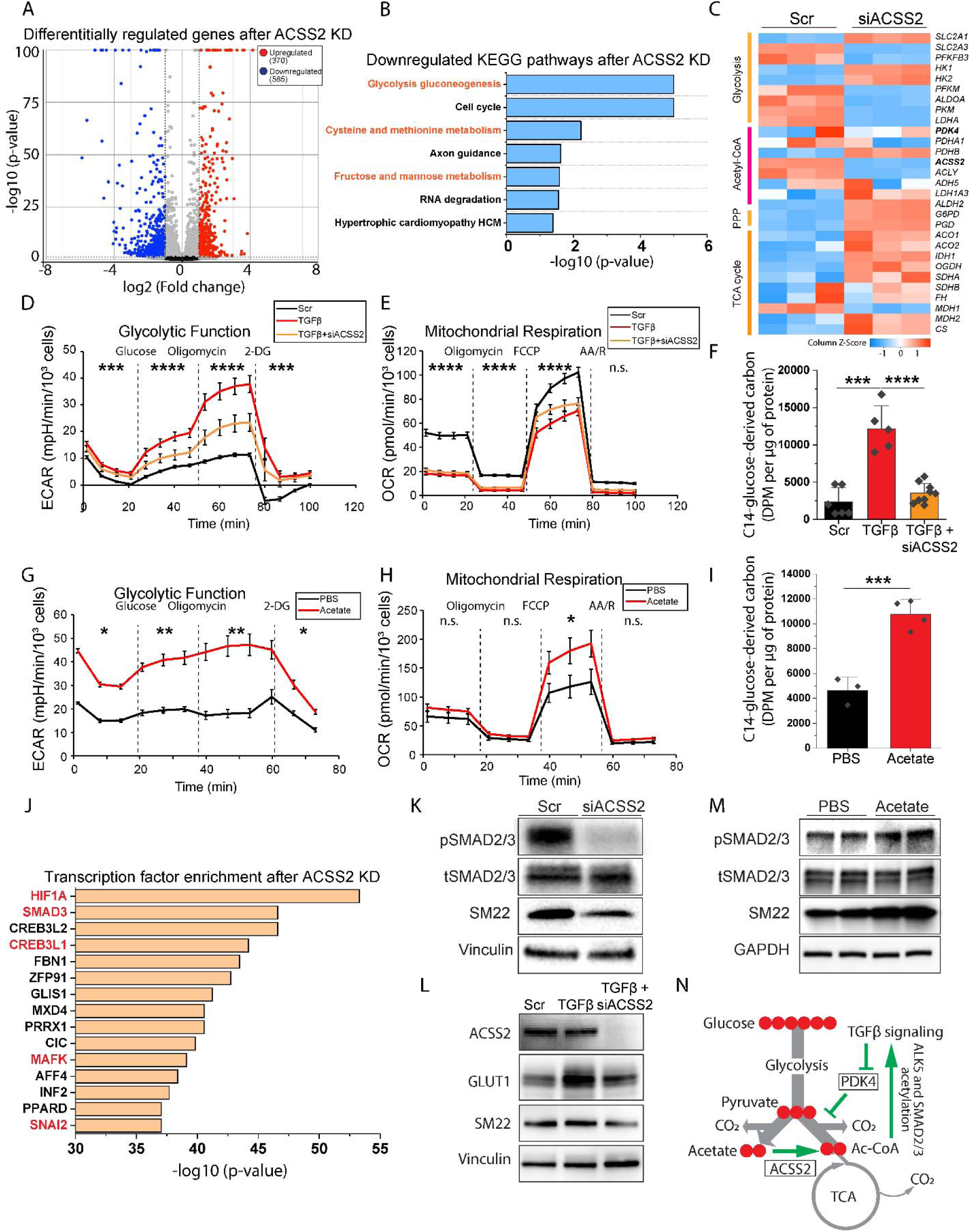
ACSS2 is essential for TGFβ activation in HASMCs. (**A-C**) Bulk RNA-seq analysis of HASMC contractile phenotype gene expression under 20 nM siACSS2 treatment for 7 days. Volcano plot (A) of differentially regulated genes upon ACSS2 knockdown. (B) Gene ontology analyses of the downregulated KEGG pathway. Orange color highlights metabolism-related pathways. (C) Heatmap of key genes involved in glucose metabolism. (**D-E**) Seahorse analyses of glycolytic function (ECAR, D) and mitochondrial respiration (OCR, E) of HASMCs treated with TGFβ or TGFβ + ACSS2 knockdown for 7 days. (**F**) Quantification of C^14^ radioactivity in CO_2_ produced by HASMCs fed with C^14^-glucose (D-[14C(U)]-glucose) for 24 h after 7 days of TGF-β stimulation with/out ACSS2 knockdown. DPM of C^14^ radioactivity was normalized to 1 µg of protein of cell lysate. (**G-H**) Seahorse analyses of glycolytic function (ECAR, G) and mitochondrial respiration (OCR, H) of HASMCs treated with 10 mM acetate for 7 days. (**I**) Radioactivity reading of C^14^-glucose-derived CO_2_ from HASMCs treated with C^14^-glucose (D-[14C(U)]-glucose) for 24 h after 7 days of 10 nM acetate treatment. (**J**) Transcription factor enrichment analysis using all the differentially regulated mRNAs after 7-day ACSS2 KD. Red text highlighted TGFβ-regulated transcription factors. (**K**) Western blotting of SMAD2/3 phosphorylation and SM22 after 7 days of ACSS2 KD. (**L**) Protein levels of GLUT1, SMAD2/3 phosphorylation and SMC contractile markers (αSMA and SM22) of HASMCs treated with TGFβ or TGFβ + ACSS2 knockdown for 7 days. (**M**) Protein levels of key genes in TGFβ signaling (phosphorylated SMAD2/3) and SMC contractile marker (SM22) after 10 nM acetate treatment for 7 days. (**N**) Diagram for Ac-CoA-induced TGFβ signaling. Green arrows indicate the TGFβ/Ac-CoA positive feedback loop, mediated by PDK4 and ACSS2. One-way ANOVA was used to compare treatment at each metabolic phase for Seahorse analyses in (D-E and G-H).

To predict the transcription factors contributing to the reduced glycolytic activities in HASMCs after ACSS2 KD, we performed ChEA using differentially regulated genes in the bulk RNAseq. We found indicated increased activities of multiple transcription factors regulated by TGFβ among the top hits following ACSS2 KD (Fig. 4j), including HIF1α(*27*), SMAD3(*12*), CREB3L1(*28*), MAFK(*29*) and SNAI2(*30*). Western blotting of SMAD2/3 phosphorylation after ACSS2 KD validated a downregulation of pSMAD2/3 levels and expression of SMC contractile phenotype markers, αSMA and SM22 levels (Fig. 4K and supplementary Fig. 3D-E). Furthermore, ACSS2 KD abolished the elevated GLUT1, pSMAD2/3 and SM22 levels induced by TGFβ (Fig. 4L and supplementary Fig. 3F). At the same time, acetate treatment induced SMAD2/3 phosphorylation as well as αSMA and SM22 levels (Fig. 4M and supplementary Fig. 3G-H). These results identify ACSS2 as an important regulator of TGFβ signaling. It mediates TGFβ-induced increase in the Ac-CoA level, which in turn activates TGFβ signaling by increasing protein acetylation of ALK5 and SMADs 2/3. Together, this leads to preservation of the SMC contractile phenotype (Fig. 4N).

### Upregulated PDK4 in atherosclerotic aortic SMCs in vivo

To investigate the relevance of the PDK4/ACSS2 molecular pathway to the SMC biology in vivo, we studied changes in SMC-specific PDK4 expression during the development and progression of atherosclerosis, a disease where SMC dedifferentiation is thought to play an important pathogenic role. To this end, we used previously published scRNAseq data (<u>GSE141031</u>) of fate-mapped aortic SMCs from *ApoE^−/−^;Myh11^−cre^;mTmG* mice on a high fat diet (HFD).

Analysis of these data showed that the aortic SMCs increased their *Pdk4* expression starting 2 months after initiation of the HFD (Fig 5A). This was associated with a decrease in expression of contractile genes, including *Acta2* and *Myh11* after 4 months of HFD (Fig. 5B-C). There was a significant negative correlation between the expression of *Pdk4* and *Acta2* or *Myh11* (R^2^ = 0.78 and 0.77 respectively, Fig. 5D-E). This suggests that a subpopulation of dedifferentiated SMCs with high PDK4 levels arises during atherogenesis. To further confirm PDK4 involvement in the development of atherosclerosis, and to localize SMCs with most significant changes in PDK4 expression, we carried out immunocytochemical analysis of ascending aortae from 8 patients with either minimal (n = 5) or mild/moderate (n = 3) extent of atherosclerosis (Fig. 5F-G). The expression of PDK4 in αSMA^+^ cells was spatially heterogeneous with the lowest expression observed in the tunica media (Fig. 5H-K and supplementary Fig. 4A-L). In particular, mean PDK4 intensity in the αSMA positive cells is 2.6-fold higher in the fibrous cap/intima than those in the tunica media (Fig. 5H&J). A strong negative correlation (R^2^ = 0.44) of αSMA vs. PDK4 expression was observed between the fibrous cap/intima and tunica media in all 8 patients (Fig. 5L). The same strong spatial heterogeneity and inverse relationship between αSMA and PDK4 levels (R^2^ = 0.87) was observed in coronary arteries of patient with severe atherosclerosis (supplementary Fig. 4N-P).

**Figure 5.**
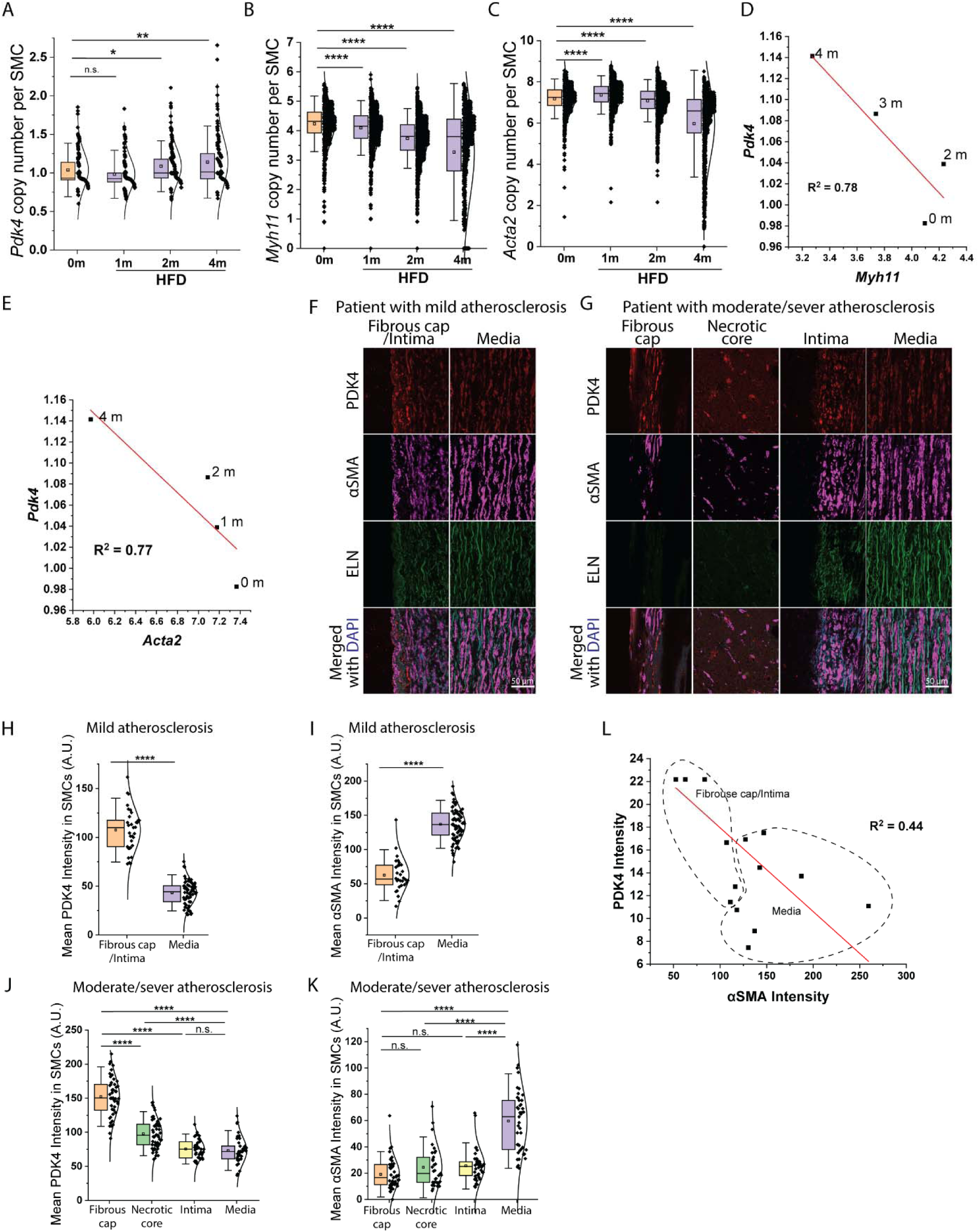
upregulated PDK4 in atherosclerotic aortic SMCs in vivo. (**A-C**) Single cell RNA-seq analyses of *Pdk4*, *Myh11* and *Acta2* expression in the aortic SMCs isolated from male *ApoE^−/−^;Myh11^−cre^;mTmG* mice under high fat diet (HFD) for 0, 1, 2 or 4 months. Each dot represents one *Myh11^−cre^*-containing GFP^+^ cells (SMCs) from one mouse in each time point. (**D-E**) Linear fitting of the average expression level of *Pdk4* to that of *Mhy11* or *Acta2* from (a-c). The time points of HFD feeding were noted next to the dots. (**F-G**) Representative images of PDK4, αSMA and DAPI staining of human patients with either mild (f) or moderate/sever (g) atherosclerosis. ELN represents the autofluorescence from elastic laminae in the tunic medium. (**H-K**) Quantification of PDK4 (H&J) and αSMA (I&K) intensity per cell from the patients with mild (H&I) and moderate/sever atherosclerosis (J&K). Each dot represents one cell. One-way ANOVA was used to test statistical significance. (**L**) Linear fitting of the normalized average expression level of PDK4 to αSMA in the fibrous cap/intima and media from all eight patients.

### SMC-specific *Pdk4* knockout ameliorates atherosclerosis in *ApoE^−/−^* mice

Given that in vitro and in vivo data imply that increased expression of SMC PDK4 induces SMC de-differentiation and promotes atherosclerosis, we investigated whether suppression of its expression would ameliorate atherosclerosis. To this end, we generated fate-mapped inducible SMC-specific *Pdk4* knockout mice (mTmG;ApoE^−/−^;Myh11CreER^T2^;*Pdk4^fl/fl^,* hereafter *Pdk4^iSMCKO^*). Activation of the Myh11CreER^T2^ by tamoxifen in mice 5–6 weeks of age resulted in a high efficiency deletion of the SMC *Pdk4* gene (Fig. 6A-C). Smooth muscle cell fate mapping showed that SMC-derived cells were effectively restricted to the tunic media after SMC *Pdk4* deletion. In line with the proof of Myh11Cre activity, immunostaining demonstrated reduced PDK4expression in SMCs in the arterial media. Whole aorta *en face* Oil Red O staining was used to assess the total atherosclerotic burden after 4 months of HFD. Compared to the ApoE^−/−^;*Pdk4^fl/fl^* mice, Pdk4^iSMCKO^ mice showed a highly significant reduction in atherosclerosis extent along the entire length of the aorta (Fig. 6D). Histologic analysis of ascending aortae demonstrated a significant reduction of the lipid-rich regions in atherosclerotic plaque of *Pdk4^iSMCKO^* mice while there were no differences in plasma cholesterol levels (Fig. 6E and supplementary Fig 5A). Taken together, these data show that the SMC PDK4 knockout reduced the extent of atherosclerosis and improved histologic compositions of the plaques.

**Figure 6.**
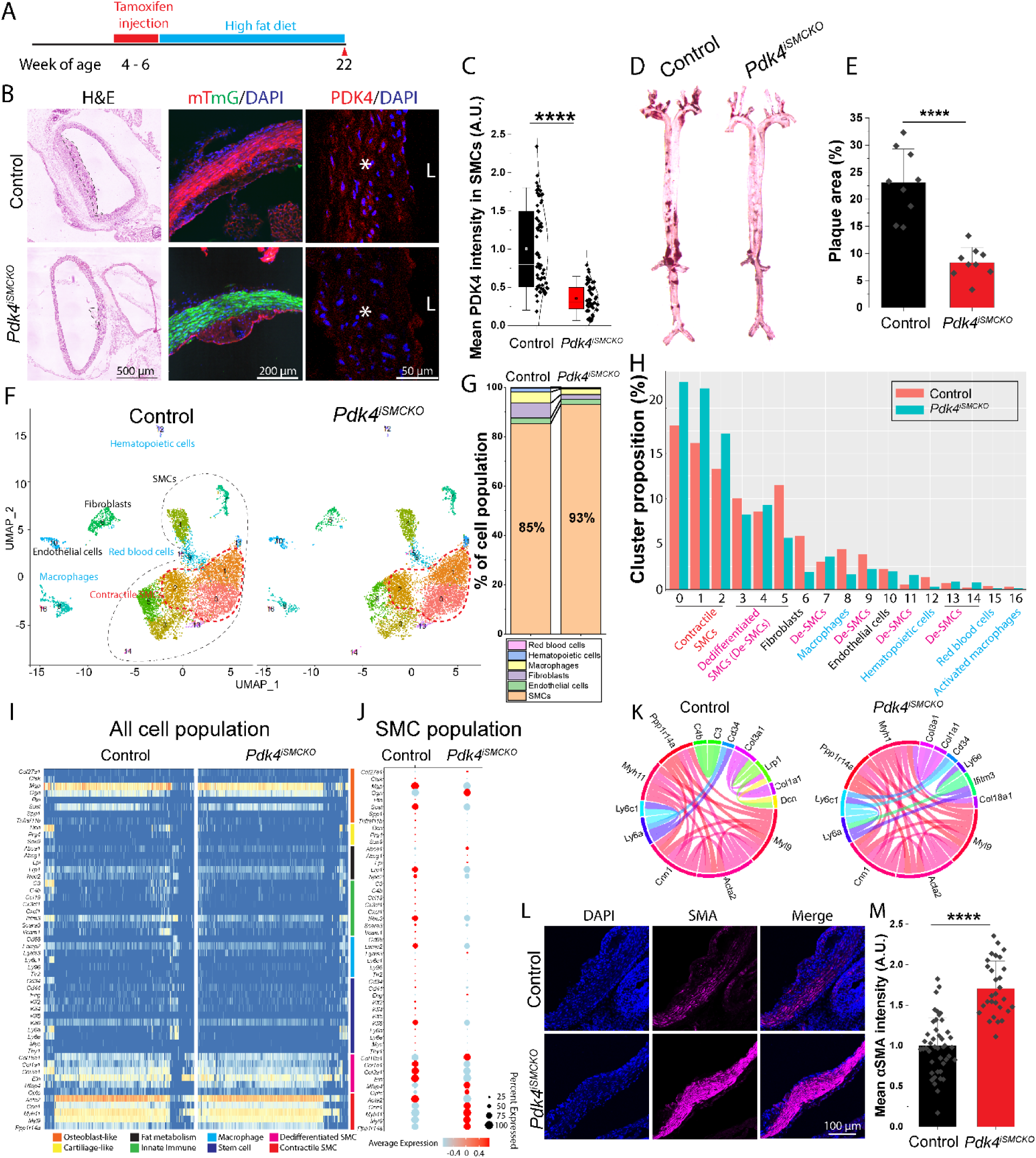
*Pdk4* knockout ameliorates atherosclerosis and maintains SMC contractile phenotype in *ApoE^−/−^*mice. (**A**) Schematic of tamoxifen injection and HFD feeding for both Control and *Pdk4^iSMCKO^* mice. (**B-C**) Representative images of H&E staining and PDK4 immunofluorescence at the ascending aortic cross-section at week 22 of age. Black dash lines outline plaque area in H&E staining. * represents the location of tunica media. “L” indicates the lumen. Quantification of mean PDK4 staining in the SMCs normalized and pooled from randomly picked 4 mice in each group were shown in (C). Each dot represents one cell. (**D-E**) Representative images (D) and quantification (E) of *en face* Oil Red O staining of whole aortic segments, including thoracic and abdominal aortae. (**F**) UMAP projections of all cells analyzed by scRNAseq for control and *Pdk4^iSMCKO^* mice. Black dash line outlines SMC population. Red dash line circles the subpopulation of contractile SMCs. (**G-H**) Population comparison among all different cell types. Note the increased percentage of SMCs in the *Pdk4^iSMCKO^* aorta in (G). (H) In-depth comparison of all the subpopulation of each cell types. Contractile SMC subgroups were marked red. Dedifferentiated SMCs (De-SMCs) were in pink. Blood cells were in blue. (**J**) Heatmap representation of scRNA gene expression for specific cell-type markers for all the isolated cells. Genes are grouped and colored by biomarker class. The colors identifying the biomarker classes are at the bottom of the heatmap. (**K-L**) Dot plot comparison (K) of differentiation markers in SMCs (*Myh11* > 0.02) between control and *Pdk4^iSMCKO^* mice. Chord plot (L) of the top 15 strongest interactions of differentiation markers in SMCs in both groups. (**M-N**) Representative images (M) and quantification (N) of αSMA staining of ascending aortic cross-sections from 4 control and 4 *Pdk4^iSMCKO^* mice. One-way ANOVA was used to test statistical significance.

### *Pdk4* knockout maintains SMC contractile phenotype in *ApoE^−/−^* mice

As the in vitro data showed that PDK4 downregulation could maintain SMC contractile phenotype via elevating TGFβ signaling, we examined this effect in *Pdk4^iSMCKO^*mice using single cell RNA sequencing (scRNAseq). Whole aortae from *Pdk4^iSMCKO^*and from control mice were collected after 4 months of the HFD and processed for scRNAseq. Cell clustering analyses showed that SMC accounted for 86% of the total cell population of the aorta in control mice, followed by fibroblasts (6%), macrophages (4.5%) and endothelial cells (2.2%). At the same time, the proportion of SMCs increased to 93% in the aorta of *Pdk4^iSMCKO^*mice while the total macrophage population decreased by 50% to 2.3% of the total (Figs. 6F-G and supplementary Fig. 5B).

Analysis of the total SMC population revealed 11 distinct clusters (Fig. 6I and Supplementary Table 1). Clusters 0,1, 2 and 4 represent the contractile SMC population with high levels of *Acta2* and *Myh11* genes expression (supplementary Figs. 5C-D). Clusters 3, 5, 7, 9, 11, 13 and 14 are the dedifferentiated HASMCs population, characterized by slightly decreased expression of contractile genes, *Acta2* and *Myh11* and increased matrix genes, *Eln* and *Col1a1*(supplementary Figs. 5E-F). Among these, Cluster 3 contains cells expressing genes characteristic of osteoblast/fibroblast-like cells and cellular inflammation (e.g. *Spp1*, *Dcn* and *Cxcl1,* supplementary Figs. 5G-I). Cluster 13 express high levels of *Klf2 and Klf4* (supplementary Fig. 5J-K). Cluster 7 is characterized by high expression of *Sox9* (supplementary Fig. 5M), a cartilage lineage marker. More in-depth cluster comparison showed that *Pdk4^iSMCKO^*mouse aorta has a higher population of contractile SMCs (Clusters 0-2 and 4, Fig. 6I), and lower population of dedifferentiated SMCs (Clusters 3, 5 and 9). In line with these data, a heatmap of SMC differentiation markers further shows that *Pdk4^iSMCKO^*aorta possesses a higher percentage of SMC contractile markers (Figs. 6J). Non-contractile SMC markers, such as SMC synthetic markers, osteoblast, stem cells and innate immune response markers, were also downregulated. A Chord plot of the top 15 most expressed markers shows a clear increase of contractile relevant genes (pink) in SMC clusters when *Pdk4* is knocked out (Fig. 6K). Immunostaining of the ascending aorta of *Pdk4^iSMCKO^* mice mouse showed increased αSMA levels (Fig. 6L-M).

### *Pdk4* knockout maintains SMC glycolysis and TGFβ signaling in *ApoE^−/−^*mice

Given phenotypic evidence of preserved contractile state of SMCs in *Pdk4^iSMCKO^* mice, we examined expression of metabolic genes after 4 months of HFD (Fig. 7A). As expected, there is a general increase in expression of glycolytic enzymes similar to that observed following *PDK4* knockdown in human SMCs (Fig. 7B). Upregulated genes include *Slc2a1 Pfkfb3, Hk1,* and *Pfkm* among others. In line with the in vitro data indicating that *PDK4* knockdown stimulates TGFβ signaling shown in Fig. 2, analysis of the scRNAseq data showed clear increases of 30 key TGFβ markers in the SMC clusters in *Pdk4^iSMCKO^*mice aorta (Fig. 7C). Immunostaining of the pSMAD2 and pSMAD3 in the aorta from *Pdk4^iSMCKO^*mice validated increased TGFβ signaling (Figs. 7D-G). These results further demonstrate that *Pdk4* knockout stimulates TGFβ signaling and glycolysis in SMC and maintains their contractile phenotype in vivo.

**Figure 7.**
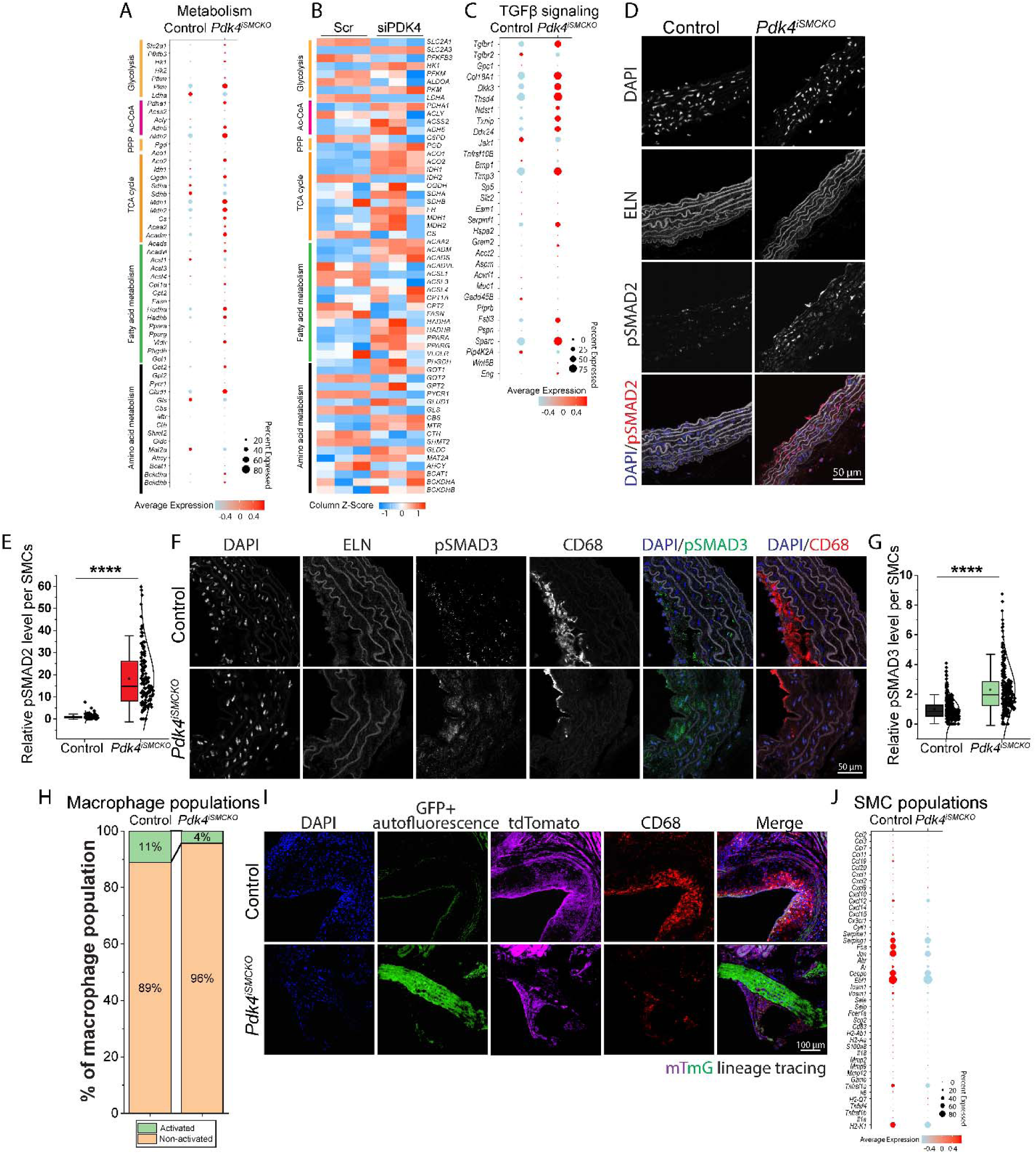
*Pdk4* knockout in SMCs reduces inflammatory responses in *ApoE^−/−^* aorta. (**A**) Dot plot comparison of key metabolic genes in SMCs (*Myh11* > 0.02) between control and *Pdk4^iSMCKO^* mice. (**B**) Heatmap of key metabolic genes in HASMCs with PDK4 KD for 7 days. (**C**) Dot plot comparison of key genes in TGFβ signaling in SMCs (*Myh11* > 0.02) between control and *Pdk4^iSMCKO^* mice. (**D-G**) Representative images of pSMAD2, pSMAD3, CD68 staining of ascending aortic cross-sections from control and *Pdk4^iSMCKO^* mice (D&F). (E&F) Quantification of relative total pSMAD2 or pSMAD3 intensity in each SMCs from 6 control mice and 5 *Pdk4^iSMCKO^* mice. One-way ANOVA was used to test statistical significance. (**H**) Population comparison of activated and non-activated macrophages. Note the reduced percentage of activated macrophage in the Pdk*4^iSMCKO^* aorta. (**I**) Representative of CD68 staining (Red) in the aortic cross-sections from mTmG-containing control and *Pdk4^iSMCKO^* mice. Green channel represents green fluorescent protein (GFP) with autofluorescence from aortic matrix. Purple channel shows the cells expression TdTomato. (**J**) Dot plot comparison of key genes in inflammation in SMCs (*Myh11* > 0.02) between control and *Pdk4^iSMCKO^*mice.

### *Pdk4* knockout in SMCs reduces inflammatory responses in *ApoE^−/−^* aorta

To verify that the reduction in atherosclerotic plaques volume in aortas of *Pdk4^iSMCKO^* mice was associated with reduced inflammation, we examined the presence and prevalence of macrophages (defined as CD68^+^ cells) in these lesions. There was a reduction in the total number of macrophages in cross-sections of the ascending aorta and the aortic root (Fig. 7F and supplementary Fig 5P) as shown by immunocytochemistry, as well as decrease in the macrophage population as determined by scRNAseq (Fig. 6F). Along with the decrease in overall macrophage numbers, there was a substantial decrease in the number of activated macrophages (defined by high level expression of *Spp1*, *Mki67* and *Gsdmd*, markers for activation, proliferation and pyroptosis respectively, supplementary Fig. 5G, N-O). Overall, the number of activated macrophages decreased 5-fold from 0.5% to 0.1% of the total cell population when *Pdk4* was knocked out in SMC. The proportion of activated macrophages in the macrophage population decreased by 64%, from 11% to 4% (Fig. 7H). This indicates that the SMC PDK4 knockout not only reduced the recruitment of macrophages into the plaque region, but also reduced the number of activated, pro-inflammatory macrophages. Macrophages in the atherosclerotic plaque are composed of blood-derived monocytes and a population of SMC that have undergone SMC-to-mesenchymal transition (SMC-MT) and began expressing macrophage markers (*4, 31*). Lineage tracing analysis shows a significant decrease in the SMC-derived macrophage population (CD68^+^, GFP^+^ and tdTomato^−^ cells) from reported 6.3-20% (*9, 32*) to 2.8% (Fig. 7I and supplementary 5Q). To explore the potential chemotactic contribution of SMC to monocyte infiltration, we studied SMC inflammatory responses. Single-cell transcriptomics showed that *Pdk4* knockout in SMCs profoundly suppresses inflammatory response as evidenced by reduced expression of genes involved in stress response (*Fos*, *Jun*, *Sperine1*, *Spering1*) and cytokine signaling (*Ccl2*, *Cxcl2*, *IL6*) (Fig. 7J). We conclude that PDK4 ablation promotes a contractile, non-inflammatory SMC phenotype, which reduces SMC-derived macrophages and limits the production of chemotactic and activating signals for infiltrated macrophages.

### SMC-specific *Pdk4* knockout slows atherosclerosis progression in *ApoE^−/−^* mice

Given the profound preventive effects of SMC-Pdk4 knockout on atherogenesis, we next examined whether the same approach would slow the progression of already established atherosclerosis. To this end, *Myh11*-CreERT2;*Pdk4*^fl/fl^;ApoE^−/−^ mice were randomized to tamoxifen-driven Cre activation (generating SMC specific *Pdk4^−/−^* ApoE mice, “*late-Pdk4^iSMCKO^*”) or DMSO treatment after 10 weeks of HFD. HFD was continued for another 6 weeks (including 2 weeks of tamoxifen/DMSO treatment), before both groups were sacrificed (Fig. 8A). The extent of atherosclerotic burden was determined using the whole aorta *en face* ORO staining (Figs. 8B&D). Additional mice were sacrificed before Cre activation to establish the baseline extent of atherosclerosis. Mice randomized to DMSO treatment demonstrated the expected progression of disease with the total aortic lesion area increasing from 8% at the start of tamoxifen treatment (Ctr-10wk) to 24% at the end of the study. At the same time, mice with the induced SMC *Pdk4* deletion (*late-Pdk4^iSMCKO^*) showed a much slower disease progression (8% to 17.5%), with plaque area at the end of the study significantly less in size than the in controls (Ctr-16wk, Fig. 8B&D). In line with the cell molecular consequences of *Pdk4^iSMCKO^* for 16 weeks, αSMA and pSMAD2 levels were increased in tunic media after L*ate-Pdk4* knockout (Fig. 8C-E).

**Figure 8.**
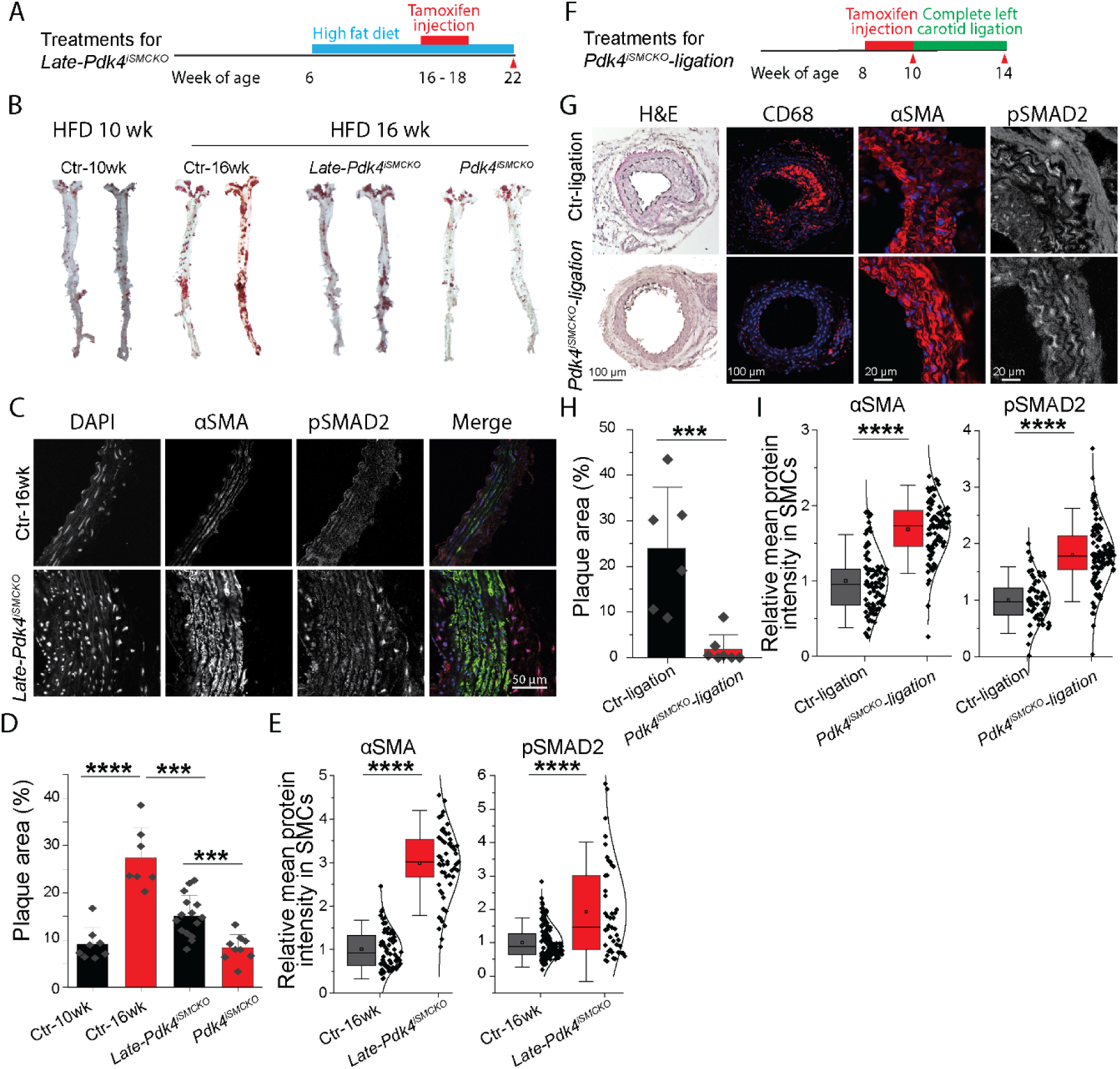
SMC-specific *Pdk4* knockout slows atherosclerosis development in *ApoE^−/−^* mice following complete carotid artery ligation. (**A**) Schematic of tamoxifen injection and HFD feeding for *Late-Pdk4^iSMCKO^* and control mice. (**B**) *en face* Oil Red O staining and quantification of whole aortic segments from mice under HFD treatments for 10 wk (Ctr-10wk) and 16 wk (Ctr-16wk). (**C**) Representative images of αSMA and pSMAD2 staining of ascending aortic cross-sections from control and *Late-Pdk4^iSMCKO^* mice (**D-E**) Quantification of plaque area in the whole aorta of each mouse (D), and relative mean αSMA and pSMAD2 intensity in each SMCs pooled from 5 Control-16wk and 4 *Late-Pdk4^iSMCKO^* mice (E). (**F**) Schematic of tamoxifen injection and left general carotid artery ligation for *Pdk4^iSMCKO^-ligation* and control mice (Ctr-ligation). (**G**) Representative images of H&E, CD68, αSMA and pSMAD2 staining of artery cross-sections from Ctr-ligation and *Pdk4^iSMCKO^-ligation* mice. Dash line outlines the plaque area in H&E staining images. (**H-I**) Quantification of plaque area percentage in each artery section (H), and relative mean αSMA and pSMAD2 intensity in each SMCs pooled from 6 Ctr-ligation and 7 *Pdk4^iSMCKO^-ligation* mice (I). One-way ANOVA was used to test statistical significance.

### SMC-specific *Pdk4* knockout reduces atherosclerosis following carotid artery ligation

Ligation of the common carotid artery is known to induce severe atherosclerosis in the ligated segment, with dedifferentiated SMCs migrating and proliferating in the ligated site (*33, 34*). Given that *Pdk4* deletion effectively maintains SMC contractile phenotype in *ApoE^−/−^* mice thereby reducing atherosclerosis burden, we set out to examine whether this would also ameliorate excessive SMC infiltration on the ligated side following a complete carotid ligation in these mice. To this end, we subjected *Pdk4^iSMCKO^* and littermate control mice without *Pdk4* gene deletion on normal chow diet to carotid ligation (Fig. 8F). Histological assessment of both mouse groups 4 weeks later showed that while control mice developed prominent atherosclerotic plaques, *Pdk4^iSMCKO^-ligation* mice were protected from as shown by the 10-fold reduction in the plaque area. Furthermore, *Pdk4^iSMCK^* was associated with significantly decreased macrophage infiltration in the plaque. Immunostaining showed increased αSMA expression (1.8-fold) and SMAD2 phosphorylation (1.7-fold) in the tunic media (Fig. 8G-H) consistent with preserved TGFβ signaling.

## DISCUSSION

The loss of SMC contractile state is thought to play a major role in the development of atherosclerosis. Our current study shows that a metabolic switch precipitated by reduced SMC TGFβ signaling input leads to a SMC phenotype switch characterized by decreased contractile gene expression and emergence of “synthetic” SMCs. The key metabolic regulator of this process is the enzyme PDK4. Low levels of PDK4 expression, induced by constitutive TGFβ signaling, allow production of cytoplasmic Ac-CoA by enzyme ACSS2. This, in turn, leads to increased acetylation of key proteins involved in the canonical TGFβ signaling, including TGFβR1 (ALK5) and SMADs 2/3, thereby stabilizing and augmenting TGFβ signaling input in a positive feedback loop. In contrast, high PDK4 expression shuts down the ACSS2-dependent Ac-CoA production and decreases expression of SMC contractile proteins. The biological and pathogenetic relevance of this PDK4-dependent SMC phenotype modulation is demonstrated both in a mouse model of atherosclerosis and in human samples. In both cases, there is an inverse relationship between the level of SMC PDK4 expression and the contractile state of SMCs. Furthermore, SMC-specific knockout of *Pdk4* both reduced the development of atherosclerosis under either HFD or complete carotid ligation and slowed atherosclerosis progression in established disease under HFD. These findings thus uncover a novel metabolic node of control in SMC differentiation and contribution to atherosclerosis (Fig. 9).

**Figure 9.**
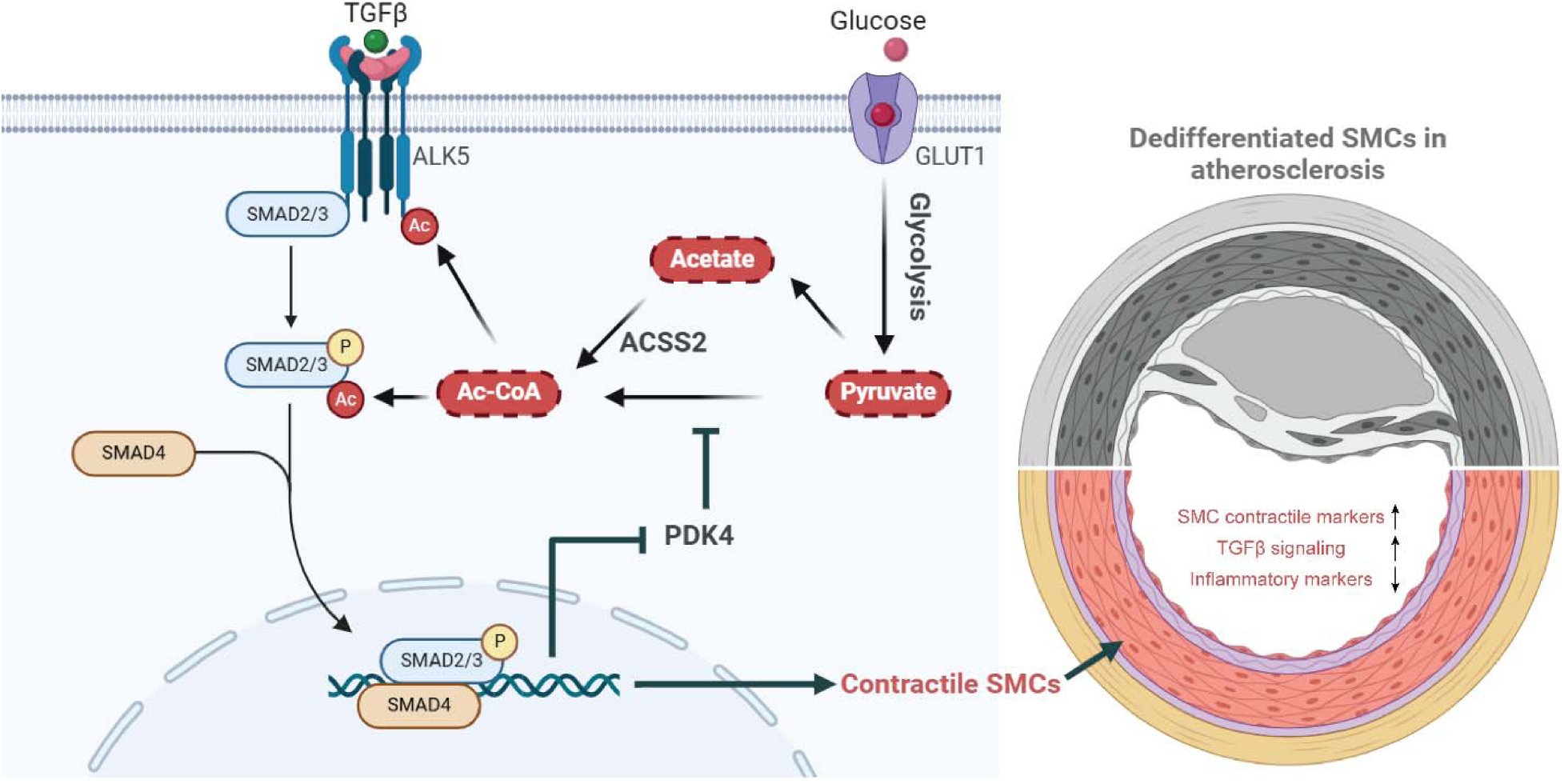
TGFβ-mediated suppression of PDK4 in SMC attenuates atherosclerosis via an Ac-CoA-dependent positive feedback loop involving SMAD2/3 acetylation.

PDK4 is a member of the PDK family of kinases and is the principal PDK expressed in SMCs. As all PDKs, PDK4 phosphorylates the PDH complex thereby inhibiting its enzymatic activity (*35*). In the absence of PDK-mediated phosphorylation, the PDH complex carries out oxidative decarboxylation of pyruvate leading to generation of Ac-CoA (that is used to drive the TCA cycle) and to produce NADH, which is used in oxidative phosphorylation to generate ATP (*36*). Two principal regulators of PDK4 expression are the transcription factor FOXO1/3 and the TGFβ-activated SMAD2/3/4. While FOXO1/3 binding to PDK4 regulatory elements increases its expression (*35*), binding of the SMAD complex decreases it (*8*). TGFβ does not directly inhibit FOXOs activities. Thus, the decreased PDK4 in SMCs by TGFβ treatment is mainly mediated by TGFβ-activated SMAD2/3/4. In addition to its classical role in driving the TCA cycle, PDH has the ability to directly convert pyruvate to acetate that can then be converted to Ac-CoA by the enzyme ACSS2 (*22*). Our data indicates that this pathway accounts for 15% of intracellular acetate and over 50% of Ac-CoA generation in SMCs when PDH is activated by suppression of PDK4. Furthermore, this is the principal pathway regulating acetylation of key regulators of TGFβ signaling, SMAD2/3, and ALK5. A higher percentage of glucose-derived Ac-CoA than glucose-derived acetate indicates that glucose-derived Ac-CoA production is from both PDH-mediated pyruvate conversion and ACSS2-mediated acetate conversion. In summary, it appears that TGFβ-mediated suppression of PDK4 leads to dis-inhibition of PDH and its non-canonical production of acetate, in turn acetylating and activating various components of TGFβ signaling, in a positive-feedback loop. It is worth noting that adding excessive exogenous acetate to the medium mimics the effect of increased glycolysis and protein acetylation, but the total oxygen consumption is also increased. It implies that exogenous acetate has some additional effect on the TCA cycle or oxidative phosphorylation. It may be that TGFβ or siPDK4-induced, i.e. PDH-derived, acetate is uniquely channeled to ACSS2 and thus cytosolic ac-CoA production, while excess exogenous acetate also has access to ACSS1 in the mitochondria to generate mitochondrial Ac-CoA, and thus feed the TCA cycle. In the context of TGFβ signaling, the cytosolic production of Ac-CoA is the important part.

In agreement with the proposed critical role of PDK4 in control of SMC homeostasis, its expression was the lowest in the tunica media of human coronary arteries where most highly contractile SMCs are located and highest in the intima and fibrous cap where most of the “synthetic” SMCs are found. Analysis of coronary arteries from patients with various severities of atherosclerosis demonstrated an inverse relationship between the extent of disease and the level of PDK4 expression. This was further confirmed by time-course studies in a mouse atherosclerosis model that show a progressive increase in PDK4 expression during development of atherosclerotic plaques. Elevated PDK4 in aortic SMCs is also associated with other cardiovascular diseases, including calcified vessels characterized by SMC dedifferentiated to osteoblast-like phenotype (*37*). Systemic knockout of *Pdk4* ameliorates vessel calcification in vitamin D3-induced mice (*37*). Intragastric administration of DCA ameliorates vessel calcification in vitamin D3-induced rats, by reducing the dedifferentiation of SMCs into osteoblast-like phenotype (*38*). These results are in line with our scRNAseq data that show decreased SMC dedifferentiated markers for osteoblast-like cells after SMC-specific *Pdk4* knockout. Taken together, these data establish a direct link between the smooth muscle PDK4 expression, SMC contractile state, atherosclerosis, and aortic homeostasis. Furthermore, these results agree with previous reports of anti-atherosclerotic effect of systemic treatment with dichloroacetate (DCA), a non-specific inhibitor of PDKs (*39*).

Another important insight from this study is the important role of SMCs in the progression of atherosclerosis. SMC-specific *Pdk4* knockout both reduced the extent of atherosclerosis development and also slowed down progression of established atherosclerosis. In both cases the likely mechanism is the decrease in the number of proliferative “synthetic” SMCs, as well as a reduction in SMC-to-mesenchymal transition (SMC-MT). The latter effect decreases the extent of SMC dedifferentiation into other cell types, including macrophage-, stem cell-, and osteoblast-like cells among other mesenchymal cell types etc. (Fig. 6I&J) (*9, 32*). Consistent with published data from lineage tracing studies that show incorporation of dedifferentiated SMCs in the plaque (*9, 31, 32, 40*), our lineage tracing results show a reduction in the number of SMC-derived macrophages in the plaque from reported 6.3-30% (*9, 32, 41*) (Fig. 7I and supplementary Fig. 5Q). This both underlies the plasticity of SMCs in these processes and points to the benefit of maintaining normal contractile SMC state.

Comparing this study and our recent work in endothelial cells (*8*), it is apparent that the metabolic circuitry controlling TGFβ signaling is very similar in SMCs and endothelium cells. In both cases, a TGFβ-mediated reduction in PDK4 and the consequent ACSS2-dependent production of Ac-CoA increases TGFβ signaling in a positive feedback loop. Furthermore, in both cell types this was associated with induction of the smooth muscle cell contractile program. The key difference, however, is that induction of the SMC contractile program in ECs induces the loss of endothelial fate gene expression, driving EndMT, a highly disease-promoting event (*5, 13*). In contrast, activation of the SMC contractile program in SMCs tends to stabilize these cells and in general suppresses vascular disease The similarity of TGFβ metabolic controls between these two cells may reflect their common developmental origin, while differences in ultimate phenotypic effect may reflect subsequent specialization into distinct cell types. This shows that the same metabolic pathway may have very different effects in different cell types. This rules out clinical use of systemic TGFβ inhibitors or stimulators as beneficial effects in one cell type would be negated by deleterious effects in another cell type. Thus, any form of TGFβ therapy would have to be cell type specific.

Our understanding of atherosclerosis has evolved significantly with much of the recent work focused on vascular inflammation and its immune modulation (*7, 9, 32, 42, 43*). Our study aligns with this paradigm, demonstrating that metabolic control of TGFβ signaling plays a critical role in maintenance of SMC contractile state and prevention of SMC-MT. This exerts an anti-inflammatory effect in atherosclerosis plaques, which is independent of changes in cholesterol levels in the blood (supplementary Fig. 5A).

An important aspect of TGFβ signaling is its cell type-specific nature. While pro-atherosclerotic role of TGFβ signaling in the endothelium and its anti-atherosclerotic effects in SMCs have previously been reported, (*8, 9*) one surprising finding in this study is that the underlying metabolic control is very similar. This may limit the appeal of PDK4 as a therapeutic target for atherosclerosis therapies. Another limitation of this study is the usage of a Y-chromosome located *Myh11*CreER^T2^ thus limiting knockouts to male mice. However, similar mechanisms are expected in female mice as there are no reported gender differences between male and female SMC metabolism. Moreover, HASMC used for in vitro studies and atherosclerotic blocks were obtained from both male and female patients, showing the same PDK4 changes and function in biological replicates from both sexes. Furthermore, a recent analysis of human smooth muscle scRNASeq data from both male and female patients show a very similar *PDK4* expression patterns (*44*).

In summary, our work unveils new metabolic underpinnings of TGFβ-mediated regulation of SMC differentiation, identifies novel targets including PDK4 for potential therapeutic development for vascular disease, and highlights the remarkable phenotypic differences that arise from similar molecular pathways in different cell types.

## MATERIALS AND METHODS

### Sex as a biological variant

Human aorta samples from both females and males were included and randomized in this research. As *Myh11*CreER^T2^ is located on the Y chromosome, only male mice can be used in this atherosclerosis study. However, similar mechanisms are expected in female mice as there are no reported gender differences between male and female smooth muscle cell metabolism.

### Immunostaining of human samples

Ascending aortas from eight organ donors with mild (five) or moderate/severe (three) atherosclerosis, and coronary arteries from 1 patient were collected by the Tellides lab. Research protocols to specimen from surgical patients were approved by the Yale Institutional Review Board with a waiver for consent (IRB # 2000020632). Research protocols for deceased organ donors were approved by the Yale Institutional Review Board with waiver of consent (IRB # 2000020632) and by the New England Organ Bank with written consent for research from the next-of-kin. All procedures were in accordance with federal and institutional guidelines. Specimen were fixed in formalin and embedded in paraffin. Blocks were sectioned at 8-µm intervals using a paraffin microtome, performed by Yale’s Research Histology Laboratory. For antigen retrieval and rehydration, paraffin sections were dewaxed in xylene, boiled at 98°C-heated 10mM Citric Acid with 0.05% Tween 20 (pH = 6) for 10 minutes. Followed by 3 times Tris-buffered saline (TBS) washes. Tissue sections were blocked by 10% BSA with horse serum (1:50) in TBS at room temperature for 1 hour. Primary antibodies were diluted in blocking solution and incubated with tissue sections overnight at 4 °C in a humidified chamber. Primary antibodies used for human paraffin section staining include anti-αSMA with cy3 fluorophore conjugated (1:1000, C6198, Millipore) and PDK4 (1:1000, 65575s, Proteintech). Sections were washed three times with TBS, incubated with donkey anti-rabbit Alexa Fluor 647 (1:200, A31573, Invitrogen) for 1 h at room temperature, washed again three times and mounted on slides with ProLong Gold mounting reagent with DAPI (Life Technologies no. P36935). All immunofluorescence micrographs were acquired using Leica SP8 microscopes. Staining intensities were quantified using imageJ as previously described (*45*). Briefly, in immunofluorescence (both direct and indirect), the intensity of protein labeling correlates with the number of epitopes bound by the primary antibody. Thus, total pixel intensity of each cell area quantified by ImageJ was used as a relative measure of the total target protein amount.

### Mice generation and tamoxifen treatment

All mouse experimental procedures used in this study were approved by the Yale University Institutional Animal Care and Use committee (protocol 2023-11231). Mice were housed in the animal facilities at Yale University, with light on 12 hrs cycles in a humidity- and temperature-controlled environment with no pathogenic microorganisms. *Myh11*CreER^T2^ mice were shared by the Tellides lab at Yale University. *Pdk4 ^fl/fl^* mice (Cyagen) (*46*) were generously provided by Dr. In-Kyu Lee from Kyungpook National University, Korea. *ApoE^−/−^* mice are from Jackson Lab (B6.129P2-Apoetm1Unc/J, Stock No. 002052). All of them are C57BL/6 background. To generate *ApoE^−/−^*;*Myh11*CreER^T2^;*Pdk4^fl/fl^*mice, we crossed these three stains, with genotyping performed by Transnetyx. Cre-Lox recombination was induced by intraperitoneal tamoxifen injection (T5648, Sigma) at 1.5 mg/day for 8 days. Tamoxifen treatment is known to reduce low density lipoprotein in the blood (*47*), which could ameliorate atherosclerosis (*48*). To eliminate the reagent artifact, mice without Cre (*ApoE^−/−^*;*Pdk4^fl/fl^*) treated with tamoxifen were used as controls by crossing male *ApoE^−/−^* mice with female experimental mice, without Y-chromosome. Mice with correct genotypes were selected using Transnetyx service (Cordova, TN) applying their proprietary real-time PCR technology.

### Atherosclerotic plaque inducements

We used two distinguished mouse models to induce vascular plaque formation: high fat diet-induced model and carotid ligated model. For the first model, mice were placed on a high fat diet (40 kcal% Fat, 1.25% Cholesterol, 0% Cholic Acid) for 16 weeks (D12108, Research Diets) starting from the age of 8-week. The common carotid ligation model was described previously (*33*). Briefly, animals were given preemptive analgesia buprenorphine XR (Ethiqa XRTM 3.25 mg/kg, SQ) and then anesthetized by an intraperitoneal injection of a mix of ketamine and xylazine (100 mg/kg and 10 mg/kg, respectively). The anterior neck incision site was infiltrated with local anesthetic lidocaine (6mg/kg). The left common carotid artery and branches were exposed through surgical cut down and ligated completely with 10-0 monofilament suture near the carotid bifurcation. The skin incision was closed using 6-0 Nylon suture and removed within 7-10 days. The right carotid artery was exposed and looped without ligation used as control. Animals were sacrificed 4 weeks after ligation for analyses of histology, immunostaining and morphometry

### Single cell RNAseq (scRNAseq) sample preparation

The mice were euthanized with isoflurane, followed by immediate perfusion with 20 mL cold phosphate-buffered saline (PBS). Isolated fresh aortae were opened longitudinally and cut into 3 mm fragments, followed by cold digestion for 3 h at 4°C. Digesting cocktail is composed of .5 mg/mL collagenase A (10103578001, Roche), 0.5 mg/mL elastase (LS002294, Worthington Biochemical), 10 mg/mL Bacillus licheniformis protease (P5380, Sigma-Aldrich), 10 mg/mL Dispase II (D4693, Sigma-Aldrich), 125 U/mL DNase I (DN25, Sigma-Aldrich) and 0.2% BSA in DMEM. Digested aortae were spined at 350 g for 5 min at 4°C, washed twice with 1% BSA in PBS, and stained with viability dye on ice for 20 min (1:2000, Thermo Fisher, 65-0865-14). Suspended cells were sorted for viable cells using flow cytometry, and collected in 0.04% BSA-PBS.

### Droplet-based scRNAseq library preparation and data analyses

10,000 cells were loaded for scRNA library preparation using the Chromium Single Cell Platform (10× Genomics) per manufacturer’s instruction. Briefly, cells were partitioned into Gel Beads in Emulsion in the Chromium system (Chromium Single Cell 3’ Library & Gel Bead Kit v4) at Yale Center for Genome Analysis, followed by cell lysis and barcoded reverse transcription of RNA, cDNA amplification and shearing, and 5’ adaptor and sample index attachment. Final scRNAseq libraries were sequenced on an Illumina NextSeq 500 using HiSeq paired-end, with read length of 150 base pairs. R package Seurat v4.4 was used for data processing, including cell filtration, normalization, principal component analysis, variable genes finding, clustering analysis, and Uniform Manifold Approximation and Projection (UMAP) dimensional reduction. To visualize the data, FeaturePlot, Vlnplot, DotPlot, chordDiagram and Heatmap were performed. Raw data were submitted to NCBI database, with approved GEO accession numbers to be <u>GSE288834</u>.

### Analysis of atherosclerotic lesions

After cardiac puncture perfusion with 20 mL PBS, the aortae were fixed with 10 mL 4% paraformaldehyde (PFA). Whole aortas, including the thoracic and abdominal segments were dissected, and fixed with 4% PFA overnight at 4°C. The entire aorta was then vertically opened with scissors, flat-mounted and rinsed three times with 60% isopropyl alcohol and stained with 0.6% Oil Red O (O0625, Millipore) at room temperature for 2 hrs. The aortae were then washed again with 60% isopropyl alcohol, followed by five times dH_2_O washes. Whole aorta images were captured with Nikon Digital Sight DS-Fi1c camera, and the surface lesion area was quantified with ImageJ software.

To measure lesions in the ascending, aortic root and carotid arteries, the fixed heart and the proximal aorta were obtained. Then they were dehydrated in 30% sucrose saline buffer at 4°C overnight before OCT embedding. OCT blocks were cross sectioned at 8 μm intervals using a Leica cryostat. Cross sections for ascending aorta were obtained 1 mm downstream of the aortic root. Those for carotid arteries were obtained 0.3 mm upstream of the ligated spot. Images of aortic roots were captured with Nikon Digital Sight DS-Fi1c camera, and the lesion area was quantified with ImageJ software. Area of plaque were measured directly using imageJ. As the artery lumen is not entirely filled with solid subtract, its area will change depending on its shape. Nevertherless, its perimeter will not change. Thus, the artery lumen area as a circle were quantified indirectly by measuring the perimeter: lumen area = π × (perimeter /2π), as previously described (*49*).

### Immunostaining of mouse tissues and quantification

All mouse tissues were frozen embedded as described in last paragraph. Before staining, slides were fixed in acetone for 10 min at −20 °C, and rehydrated in PBS at room temperature. After washing three times with TBS, tissue sections were blocked and incubated with primary antibodies as described previously. Primary antibodies used include anti-αSMA (1:1000, C6198, Millipore), anti-pSMAD2 (1:1000, ab3849-1, Sigma-Aldrich), anti-pSMAD3(1:1000, ab52903, Abcam), anti-CD68 (1:1000, MCA1957, Bio-Rad). Sections were washed three times with TBS, incubated with appropriate Alexa Fluor 647 or Alexa Fluor 594 conjugated secondary antibodies diluted 1:200 in blocking buffer for 1 h at room temperature. Three times TBS washes were performed before mounting with ProLong Gold mounting reagent with DAPI (Life Technologies no. P36935). All immunofluorescence micrographs were acquired using Leica SP8 microscopes, with same laser intensity applied to compare different conditions. Staining intensities were quantified using ImageJ as previously described (*45*).

### Cell culture and treatments

Primary Human Aortic smooth muscle cells (HASMCs) from two donors were purchased from Promocell (C-12533) and Lonza (CC-2571) respectively. The HASMCs were cultured in Smooth Muscle Cell Growth Medium-2 (SmGM-2, CC-3181, Lonza), supplemented with BulletKit (CC-4149, Lonza), 100 µg/mL penicillin and 100 µg/mL streptomycin. Cell culture dishes were maintained in a humidified 37 °C incubator with 5% CO_2_.

For TGFβ or acetate treatments, 10 ng/ml TGFβ2 or 10 mM acetate were added into the complete SmGM-2 growth media for 7 days with media changes every other day. For RNAi experiments, 20 µM of siRNA for PDK4 (s10263, Thermo Fisher Scientific), ACSS2 (E-010396-00-0005, Horizon Discovery Biosciences), SLC2A1 (L-007509-02-0005, Horizon Discovery Biosciences) or SMAD4 (s8403, Thermo Fisher Scientific) were transfected with lipofectamine RNAimax (13778075, Invitrogen) in OptiMeM (31985062, Invitrogen) overnight, followed by complete medium. Transfections were performed every 72 h, with scrambled siRNA as control (4457287, Thermo Fisher Scientific). For overexpression experiments, adenovirus of PDK4 (1:2000, 363230510100, Applied Biological Materials) and ACLY (1:2000, 11175051-HA, Applied Biological Materials) were applied to HASMC culture for 3 days. Same dilution of adeno CMV null adenovirus (000047A, Applied Biological Materials) was used as control. The cells were maintained between 70-90% confluence throughout the experiments. Inhibitors used in this paper include 10 μM ALK5 inhibitor (SB525334, MCE, HY-12043), 50 μM PDK4 inhibitor (PDK4-IN-1, MCE, HY-135654) and 50 μM PDK inhibitor (DCA, MCE, HY-Y0455A).

### ATAC-RNAseq sample preparation and sequencing analyses

ATAC-RNAseq was performed at Yale Center for Genome Analysis. 100,000 treated HASMCs were trypsinized and cryopreserved in 90% fetal bovine serum + 10% dimethyl sulfoxide. Fastqc was used to check the quality of the data and identify the adapters. Trimmomatic was used to cut the adapters. The trimmed data then was aligned to human reference genome, hg38. Quality control of mapped data, including sort the data, make duplicate reads, and removing ENCODE blacklist were performed using the samtools and bedtools. MACS2 was used to call the peaks. Mapped data was converted to bigwig files using deepTools, and visualized by Integrative Genomics Viewer. Raw data were submitted to NCBI database, with approved GEO accession numbers to be <u>GSE288977</u>.

### Total RNA isolation, Bulk RNA-seq analysis and qPCR analysis

Extraction of total RNA was performed using the RNeasy Mini Kit (14734, Qiagen), according to the manufacturer’s instructions. Novaseq sequencings with 25 million reads per sample were performed to detect the whole transcriptome. RNA-seq data was aligned to the reference human genome (Hg19) using STAR aligner on PartekFlow platform. Strict paired-end compatibility was applied for gene expression quantification. Read counts were normalized using the trimmed mean of M-values method. Differentially expressed genes with a false discovery corrected p ≤ 0.05 were used for further analysis (heatmaps, volcano plots and functional enrichment). Bulk RNA-seq datasets were uploaded into NCBI database, with approved GEO accession numbers to be <u>GSE288577</u> and <u>GSE288831</u>. To confirm the different expression of genes involved in SMC differentiation and Glucose/Ac-CoA metabolism, first-strand cDNA was synthesized using iScript cDNA synthesis lit (1708890, Bio-RAD), and gene expressions were analyzed using iQ™ SYBR® Green Supermix (1708880, Bio-RAD). β*-ACTIN* was used as house-keeping gene. Primers are listed in Supplementary Table 2.

### Liquid chromatography-mass spectrometry (LC-MS) metabolomics analysis

To trace glucose-derived metabolites, cultured HASMCs were switched to DMEM media without glucose or pyruvate (11966025, Gibco), supplemented with 5% dialyzed serum (A3382001, Gibco) and labeled glucose (10 mM U-^13^C6-Glucose, CLM-1396-1, Cambridge Isotope). After 1-day incubation in a CO_2_ incubator, metabolites were extracted from 1.5 × 10^6^ cells per well using 3 mL of ice-cold 80% MeOH for 15 min. Cells were collected by rubber policemen and transferred to three 1.5ml Eppendorf tubes on dry ice. The samples were homogenized by vortexing for 10 seconds and centrifuged at 16,000 g for 15 min at 4 °C. The supernatant extract from the same sample was pooled into one 15 ml conical tube on dry ice. Extracts were dried and concentrated by 10-fold with nitrogen blow. The samples were then centrifuged at 16,000 g for 10 min at 4°C, and 40 μL of supernatant was used for LC-MS run.

Cell extracts were loaded and analyzed using quadrupole-orbitrap mass spectrometer (Q-Exactive Plus Hybrid Quadrupole-Orbitrap, Thermo Fisher) coupled to Vanquish UHPLC Systems (Thermo Fisher) via electrospray ionization. They were electrospray ionized via hydrophilic interaction chromatography (HILIC). LC separation was performed using Xbridge BEH amide column (2.1 mm x 150 mm, 2.5 μm particle size, 130 Å pore size; Waters) at 25 °C using a gradient of solvent A (5% acetonitrile, 20 mM ammonium acetate and 20 mM ammonium hydroxide) and solvent B (100% acetonitrile). Flow rate was 150 μL/min. The LC gradient was: 90% B (0-2 min); 75% B (3-7 min); 70% B (8-9 min); 50% B (10-12 min); 25% B at 13 min; 20% B (14-15 min); 0% B (16-20.5 min); 90% B (21-25 min). 3 μL of samples were injected with autosampler temperature at 4°C. MS data were acquired with a full-scan mode from m/z 70 to 830 and 140,000 resolution in negative ion mode. Peaks were identified with MAVEN software. Natural isotope correction was performed with AccuCor2 R code (https://github.com/wangyujue23/AccuCor2). Total ion counts refer to the sum of all unlabeled and labeled forms, in which each form is weighted by fraction carbon atoms labeled. These counts were utilized to determine fractional carbon labeling (%) by normalizing against the ion count of the total pool. Additionally, total ion counts were adjusted based on cell numbers.

### Ac-CoA quantification by LC-MS

Samples were collected and analyzed as previously described in detail by liquid chromatography- high resolution mass spectrometry (*50*). Briefly, after aspirating the media, 1.5 × 10^6^ HASMCs were lysed with 1ml of ice-cold 10% (w/v in water) trichloroacetic acid (TCA) (T6399, Sigma), transferred to 1.5 mL Eppendorf tubes, spiked with 0.1 mL of ^13^C315N1-ac-CoA internal standard prepared as previously described from yeast (*51*), then homogenized by vortexing and 12 × 0.5 s pulses with a probe tip sonicator. Protein was pelleted with 17,000 g centrifugation for 10 min at 4 °C. Solid-phase extraction was performed with the supernatant loaded to Oasis HLB 1cc (30 mg) solid phase extraction columns (Waters) previously equilibrated with 1 mL of methanol then 1 mL of water. SPE columns were then desalted with 1 mL water and eluted with 1 mL methanol containing 25 mM ammonium acetate. The purified extracts were evaporated to dryness under nitrogen, then resuspended in 55 μl 5% (w/v) 5-sulfosalicylic acid in water. Calibration curves were prepared from commercially available ac-CoA standards (Sigma Aldrich) in the same manner. 10 µL injections were analyzed on an Vanquish Duo UHPLC using a Waters HSS T3 2.1 × 100mm 3.5 µm column coupled to a Q Exactive Plus. The [M+H]+ ions of Ac-CoA was used for quantification. Data were integrated using Tracefinder 5.1 (Thermo Scientific) software. The analysts were blinded to sample identity during processing and quantification.

### Seahorse assay of oxygen consumption rate (OCR) and extracellular acidification rate (ECAR)

To investigate the metabolic changes of HASMCs, OCR and ECAR were measured with Agilent Seahorse XF96 analyzer (Agilent Technologies). In brief, HASMCs per well after 6-day treatment were transferred and seeded on Seahorse XF96 microplate (101085-004, Agilent) at 37 °C in a CO_2_ incubator overnight before the assays. 30,000 HASMCs were seeded per well. At the same time, sensor cartilage (102601-100, Agilent) was hydrated with Seahorse XF Calibrant solution (100840-000, Agilent) and incubated at 37 °C overnight without CO_2_. On the day of the experiment, HASMCs were changed into FBS-free Agilent Seahorse XF DMEM Media (pH7.4, 103575-100, Agilent) in a non-CO_2_ incubator for 1 h before the assay. For OCR measurements, Oligomycin (3 μM), FCCP (1 μM) and Antimycin/Rotenone (1.5 μM/3 μM) were sequentially injected to the medium. For ECAR measurements, glucose (5 mM), oligomycin (3 μM) and 2-DG (100 mM) were sequentially added to the cells according to the manufacturer’s protocols.

### Fluorescent tracer assay for glucose uptake

HASMCs culture medium were supplemented with 5 mM 2-(7-Nitro-2,1,3-benzoxadiazol-4-yl)- Deoxyglucose (2-NBDG, N13195, Invitrogen). After 4 h incubation, live cells were washed with PBS containing 5 mM glucose for three times and mounted in it. Images were taken using Leica SP8 microscopes for fluorescence of 488 nm wavelengths. As 2-NBDG can be uptake by glucose transporters, and cannot be further metabolized, fluorescence intensity reveals levels of glucose uptake. Method for 2-NBDG intensity quantification was described previously (*45*).

### Radioactive measurement for glucose oxidation

To chase the glucose-derived CO_2_, treated HASMCs were incubated with 5 mM radiolabeled D-[^14^C(U)]-glucose (NEC042V, Perkin Elmer) in glucose-deprived DMEM medium containing 5% dialyzed serum for 24 h. CO_2_ was released from the HASMCs and culture medium with 3M perchloric acid (1:2 v/v). The released CO_2_ was captured by filter paper soaked with hyamine. Disintegrations per minute (DPM) for radioactive CO_2_ was read by liquid scintillation counter (Perkin Elmer) in scintillation vials. Finally, the data was normalized by cell number.

### Acetylation test for ALK5 and SMAD2/3 proteins

6 hours before cell collection, 5 uM Trichostatin A (TSA) was added to HASMCs culture to inhibit deacetylase activities. Cells were lysed in ice cold RIPA buffer containing protease inhibitors and 5 uM TSA. The lysate were precleared with agarose beads conjugated with protein G, followed 12,000 g centrifuge for 5 min. Surprenant were incubated with agarose beads conjugated with anti-Acetylated lysine antibody (AAC04-Beads, Cytoskeleton) to enrich acetylated proteins at 4 °C with gentle rotation overnight. The beads compound with acetylated proteins were collected by gentle spin for 2 min at 3, 000 rpm, followed by seven times washes with ice cold RIPA buffer with 5 uM TSA. Elution was performed using 2 × Laemmli Sample buffer at room temperature for 5 min. Eluted proteins were denatured with 2 μl β-mercaptoethanol and heated in 95 °C for 5 min for western blotting.

### Western blotting

Western blotting was performed as previously described (*49*). One of the following primary antibodies at a 1:1,000 dilution in 5% BSA in TBS: Acetylated-Lysine (Cell Signaling, 9441S), ACSS2 (Cell Signaling, 3658S), ALK5 (Abcam, ab235578), αSMA (R&D Systems, IC1420A), GAPDH (Cell Signaling, 2118S), GLUT1 (Abcam, ab652), PDK4 (Proteintech, 12949-1-AP), PFKFB3 (Proteintech, 13763-1-AP), phosphor-PDH (S293) (Abcam, ab92696), pSMAD2 (Ser465/467) (Millipore, AB3849), pSMAD3 (Ser423/425) (Abcam, ab52903), SM22 (Abcam, ab14106), SMAD2/3 (Cell Signaling, 3102S) and Vinculin (Proteintech, 26520-1-ap). Chemiluminescence measurements were performed using SuperSignal West Pico Chemiluminescent Substrate (Thermo Fisher Scientific Prod #34080).

### Cell viability assay

The MTS (3- (4, 5-dimethylthiazol-2-yl)-5- (3-carboxymethoxyphenyl)-2- (4-sulfophenyl)-2H-tetrazolium) assay (Promega, G5421) was used as manufacturer instructed. 10,000 HASMCs per well after 7 days of TGFβ or PDK4 knockdown treatment were seeded in 96-well plate. Absorbance was read at 570 nm.

### Plasma lipid analysis

Before sacrifice, mice after 15-week HFD treatment were starved overnight. Blood samples were collected by cardiac puncture and mixed with EDTA (25 mM EDTA per 1 mL plasma). The samples were then centrifuged at 10,000 g at 4°C for 10 min. Serum were collected for to measurements of total and high-density lipid cholesterol levels according to the manufacturer’s instructions (FujiFilm Healthcare, 999-02601 & 997-01301).

### Statistical analysis

Sample sizes were determined by Power Analysis based on preliminary data obtained in the lab. Blinded quantification was performed to avoid bias. Statistical analyses were performed using Origin2022. The data are presented as the mean ± standard deviation from at least three independent experiments. Prior to the significance analyses, outliers were determined by the Grubbs’ test at a 95% confidence level and excluded. Two-sample statistical analyses were performed using one-way ANOVA. In relative analyses, the averages of the control groups were normalized to 1, and paired-sample t-test were performed. One-sample t-test was used in supplementary Fig 5Q. p < 0.05 (*), p < 0.01 (**), p < 0.001 (***) and p < 0.0001(****) were considered statistically significant.

## Acknowledgments

Acknowledgment of nonauthor contributions: We thank Dr. Yajaira Suarez for generously sharing the seahorse assay machine for this study.

## Funding

National Institutes of Health grant, R01 HL135582, (MS)

National Institutes of Health grant, HL167014, (MS)

Coefficient Giving, (MS)

Knut and Alice Wallenberg Foundation, KAW 2020.0057, (MS)

Marfan Foundation, Victor McKusick Fellowship, (RMZ).

## Author contributions

Study conception and design: RMZ, XZ, MAS, ZA and MS.

Acquisition of data: RMZ, JZ, NS, HB, YL and PYC.

Manuscript writing: RMZ, ZA and MS.

Materials and equipment providing: JZ, HS, GT, NS, CJ, MAS and MS.

## Data, code, and materials availability

All data and code needed to evaluate and reproduce the results in the paper are present in the paper and/or the Supplementary Materials. Raw sequencing data were submitted to NCBI database, with approved GEO accession numbers <u>GSE288834</u>, <u>GSE288977</u>, <u>GSE288577</u> and <u>GSE288831</u>. Customed codes for R package Seurat v4.4 to analyze scRNAseq are available in Dryad (<u>https://doi.org/10.5061/dryad.m0cfxppjn</u>). This study did not generate new materials.

## Supplementary Materials

**Supplementary figure 1.**
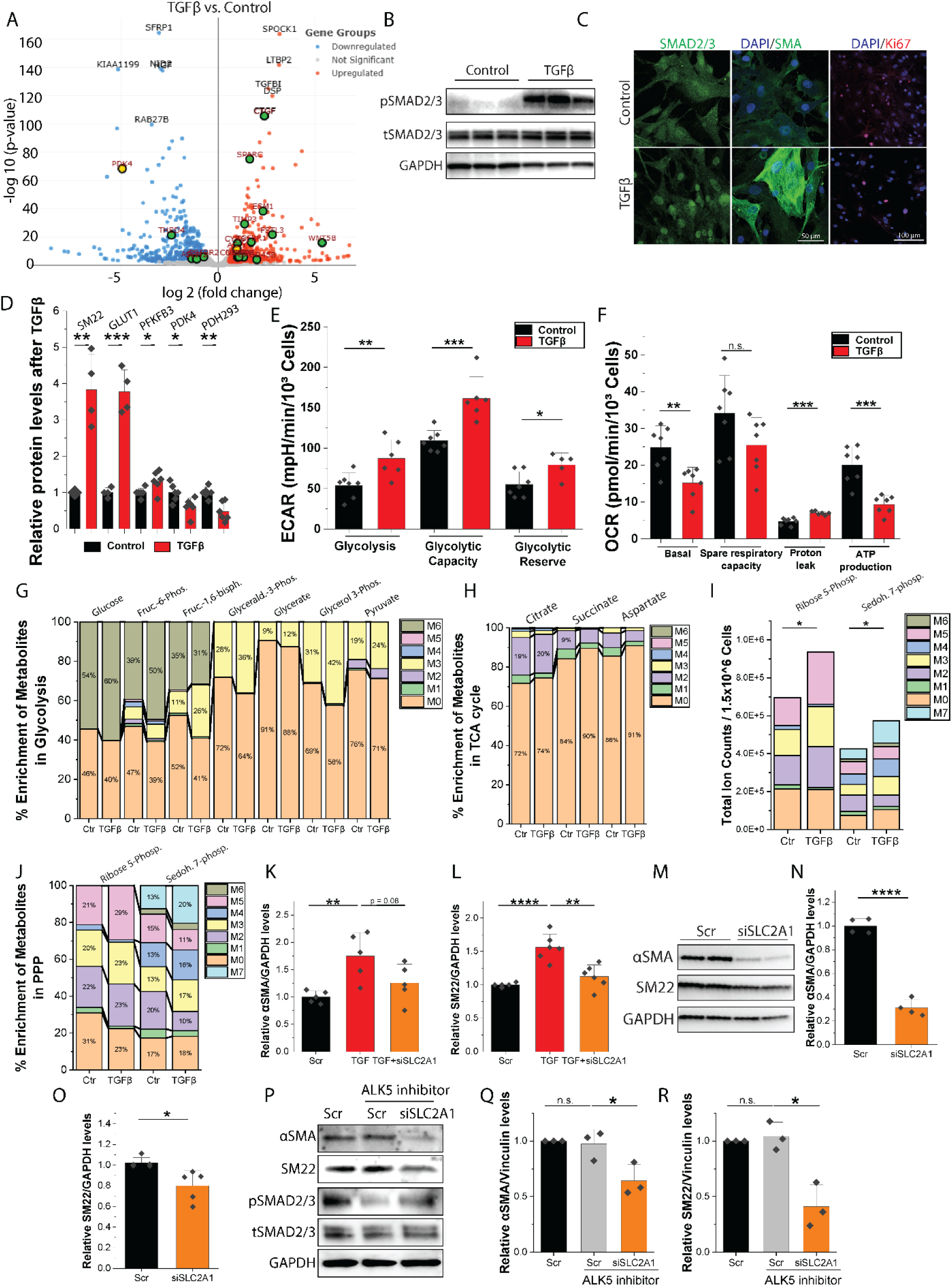
Metabolic effects of HASMC TGFβ signaling. (**A**) Volcano plot of transcriptome in TGFβ treated HASMC compared to control. Green dots represent TGFβ-targeted genes. Yellow dots represent glycolytic enzymes. Note that PDK4 is among the most downregulated genes after 7-day of TGFβ treatment. (**B**) Western blotting of TGFβ signaling (SMAD2/3 phosphorylation). (**C**) Immunostaining of SMAD2/3, αSMA and Ki67. Note that TGFβ promotes the SMAD2/3 translocation to the nucleus, and induces HASMCs to a more contractile phenotype with increased αSMA and decreased Ki67. (**D**) Quantification of SM22, GLUT1, PFKFB3, PDK4 and PDH293 after 7-day of TGFβ treatment. (**E-F**) Seahorse analyses of (E) glycolysis, glycolytic capacity and glycolytic reserve and (F) basal oxygen consumption, spare respiratory capacity, proton leak and ATP production of HASMCs after treated with 10ng/mL TGFβ for 7 days. (**G-J**) LC-MS metabolomics analysis of key metabolites in HASMCs treated with C^13^-glucose (U-13C6-glucose, 10 mM) for 24 h after 7 days of TGF-β stimulation. M# represents the number of C^13^ -labeled carbon in each metabolite. Note that the % of C^13^ labelled metabolites in glycolysis (G) and pentose phosphate pathway (PPP, H-I) were increased by TGFβ, whereas that of key metabolites in the TCA cycle(H) were not increased. (**K-L**) Quantification of αSMA and SM22 levels, normalized to GAPDH, to investigate the effects of GLUT1 KD (siSLC2A1) on HASMC contractile phenotype under 10 ng/mL TGFβ treatments for 3 days (**M-N**) Representative western blots and quantification studying the effects of GLUT1 KD (siSLC2A1) on HASMC contractile phenotype (αSMA and SM22) under normal growth medium, with baseline TGFβ level, or with ALK5 inhibitor treatment for 3 days. One-way ANOVA was used for statistical analyses in (D, E, F, I, K, L, N and O). Paired-sample t-test was used in (Q and R).

**Supplementary figure 2.**
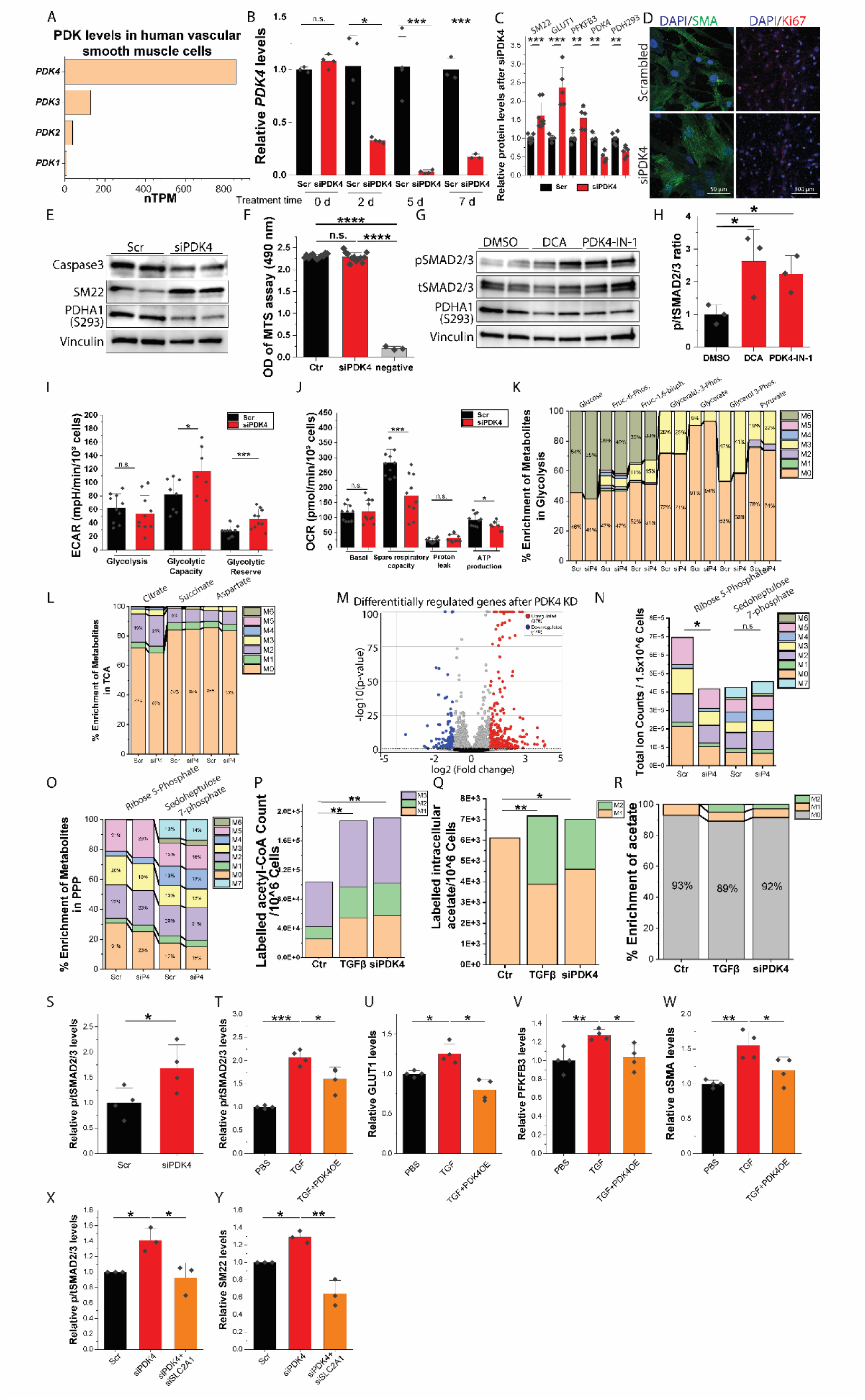
Metabolic effects of PDK4 knockdown in HASMCs. (**A**) Transcripts per million (nTPM) of four PDK isoforms in human vascular aorta. (**B**) Time course qPCR analyses of relative PDK4 levels upon siPDK4 treatment for 0, 2, 5 and 7 days. (**C**) Immunostaining of αSMA and Ki67 of HASMCs after PDK4 KD for 7 days. (**D**) Quantification of SM22, GLUT1, PFKFB3, PDK4 and PDH293 after PDK4 KD for 7 days. (**E-F**) Western blot of Caspase-3 and MTS assay in HASMC after PDK4 KD for 7 days (**G-H**) Western blot and quantification of SMAD2/3 phosphorylation after DCA or PDK4-IN-1 treatment for 3 days. (**I-J**) Seahorse analyses of (I) glycolysis, glycolytic capacity and glycolytic reserve and (J) basal oxygen consumption, spare respiratory capacity, proton leak and ATP production of HASMCs after PDK4 KD for 7 days. (**K-L**) LC-MS metabolomics analysis of key metabolites in HASMCs treated with C^13^-glucose (U-13C6-glucose, 10 mM) for 24 h after PDK4 KD for 7 days. M# represents the number of C^13^-labeled carbon in each metabolite. Note that the % of C^13^ labelled metabolites in glycolysis (K) were increased by TGFβ, whereas that of key metabolites in the TCA cycle (L) were not increased. (**M**) Volcano plot from bulk RNA-seq analysis of HASMC contractile phenotype gene expression under 20 nM siPDK4 treatment for 7 days. (**N-O**) Total ion counts (N) and percentage (O) of C^13^-labeled metabolites in PPP. HASMCs treated with C^13^-glucose (U-13C6-glucose, 10 mM) for 24 h after PDK4 KD for 7 days. (**P**) Total ion counts of C^13^-glucose derived-Ac-CoA quantification by LC-MS in HASMCs treated with TGFβ or PDK4 knockdown for 7 days. M# refers to the C^13^ number in one molecule. (**Q-R**) Total ion counts and % of C^13^ labelled intracellular acetate in HASMCs treated with TGFβ or PDK4 KD for 7 days. (Q) includes the non-labelled acetate (M0). Note that no M2 labelled acetate was observed in control group. One sample t-test was performed for M2 acetate in the TGFβ or PDK4 KD group. (**S**) Quantification of phosphorylated to total level of SMAD2/3 after PDK4 KD for 7 days. (**T-W**) Quantification of phosphorylated to total level of SMAD2/3, GLUT1, PFKFB3, αSMA levels after 3-day treatment with 10ng/mL TGFβ with PDK4 adenovirus or control virus for 3 days. (**X-Y**) Quantification phosphorylated to total level of SMAD2/3 and SM22 levels after 7 days of PDK4 KD or PDK4 and GLUT1 double KD. One-way ANOVA was used for statistical analyses of (B, D, F, H, I, J, N, P, Q, S, T, U, V and W). Paired-sample t-test was used in (X and Y).

**Supplementary figure 3.**
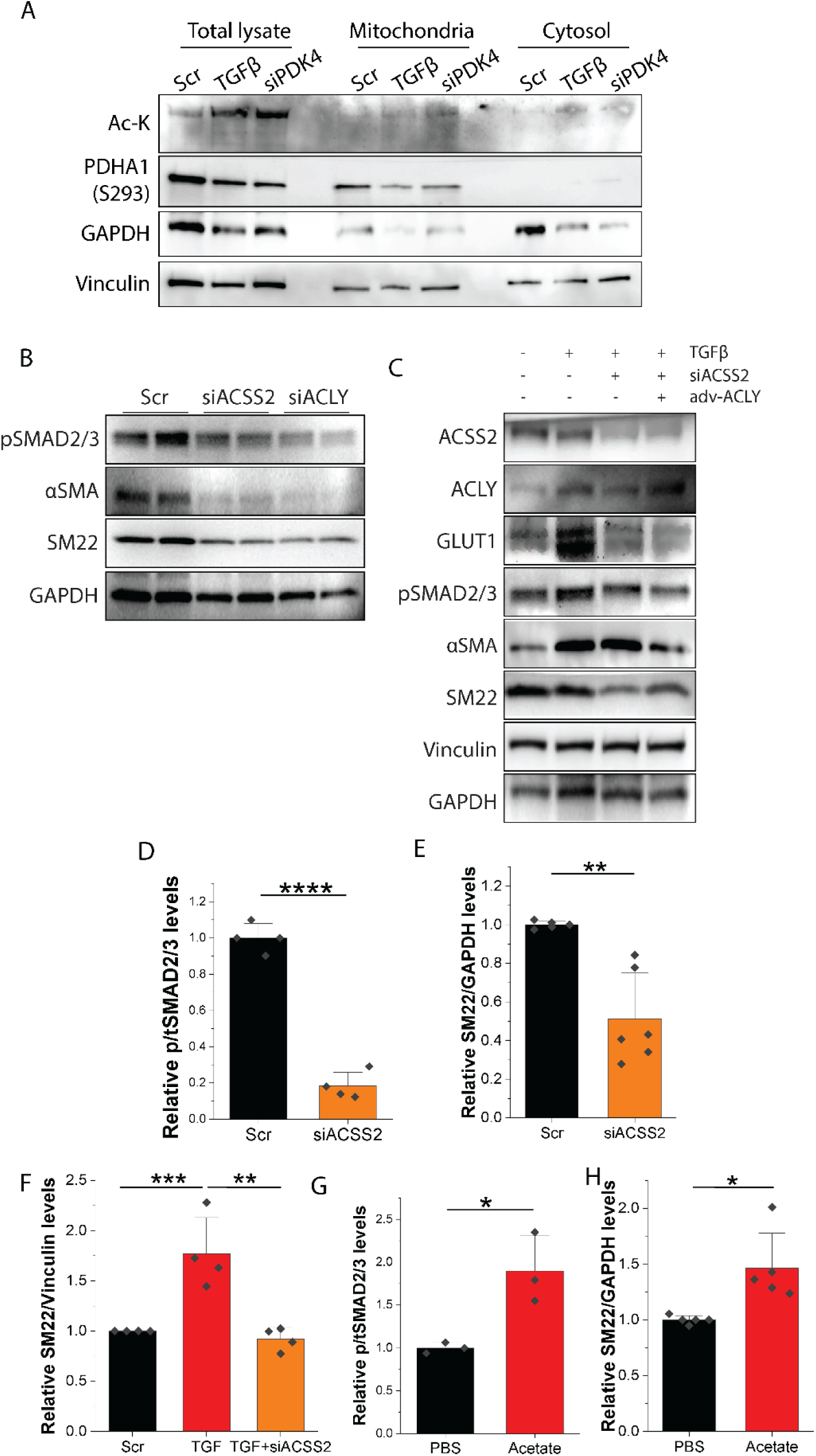
Analysis of Subcellular distribution of protein acetylation. (**A**) Western blotting of acetylated protein, using anti-acetylated lysine (Ac-K). Phosphorylated PDHA1 (S293) is used as mitochondria marker. GAPDH is mainly in the cytoplasm. (**B**) Western blotting of pSMAD2/3 and SMC contractile markers (αSMA and SM22) upon ACSS2 or ACLY knockdown for 3 days. (**C**) Rescuing experiments of adenoviral overexpressing ACLY (adv-ACLY) on TGFβ signaling (pSMAD2/3), SMC contractile phenotype (αSMA and SM22) and glucose uptake (GLUT1), after ACSS2 knockdown. (**D-H**) Quantification of phosphorylated to total ratio of SMAD2/3 and SM22 levels, normalized to GAPDH or Vinculin, to investigate the effects of ACSS2 KD and 10 mM acetate treatment on HASMC contractile phenotype. One-way ANOVA was used for statistical analyses for (D, E, G and H). Paired-sample t-test were applied to (F).

**Supplementary figure 4.**
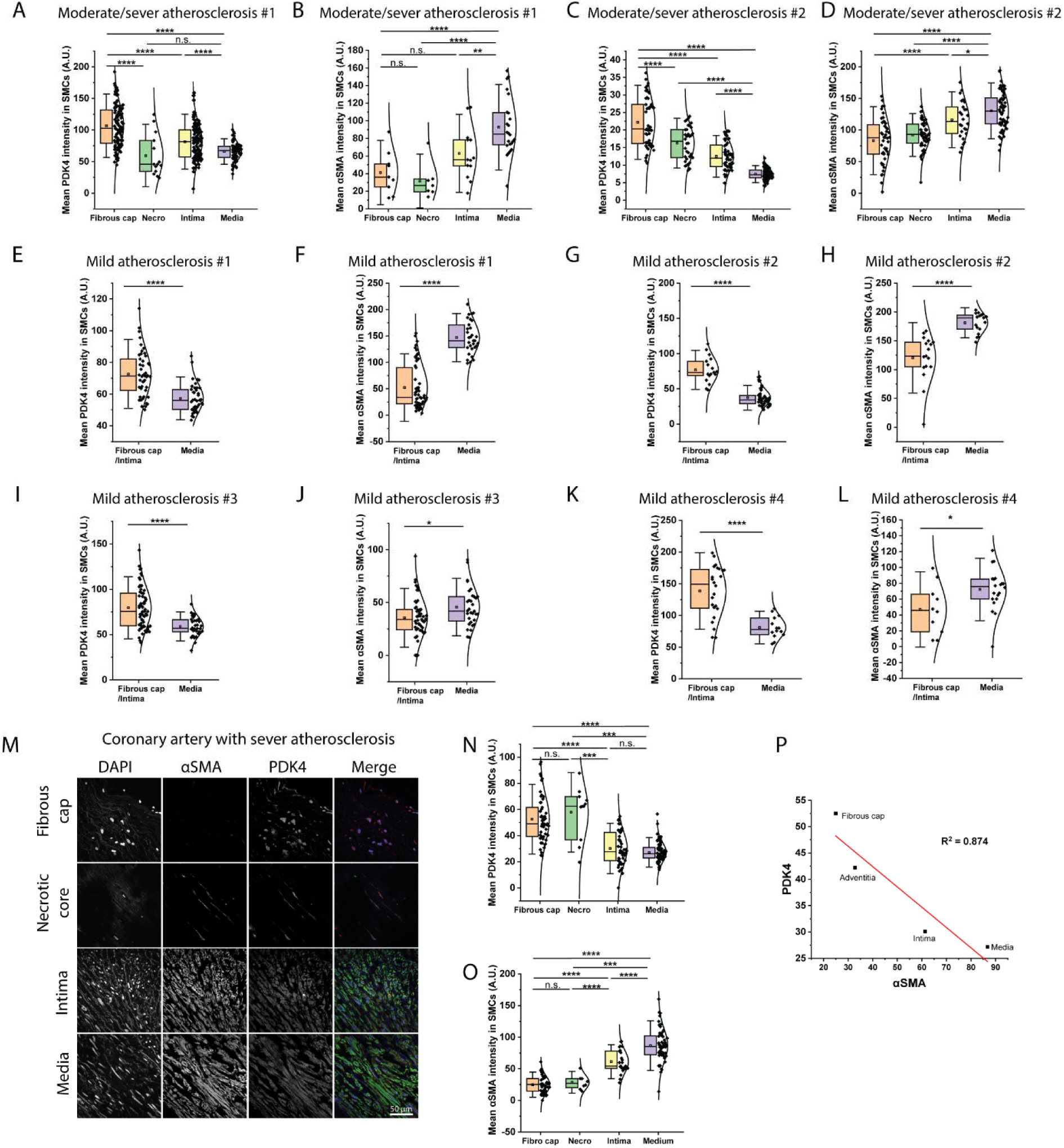
Upregulated PDK4 in atherosclerotic SMCs in human aortae and coronary arteries. (**A-L**) Quantifications of PDK4 and αSMA staining in human ascending aorta from human patients either with moderate/sever (A-D) or mild (E-L) atherosclerosis. Each dot represents one SMCs. (**M-O**) Representative images (M), quantification (N-O) of PDK4 and αSMA intensity of coronary artery from a patient with sever atherosclerosis. (**P**) Linear fitting comparing mean PDK4 and αSMA intensity from different locations in coronary artery. One-way ANOVA was used for statistical analyses

**Supplementary figure 5.**
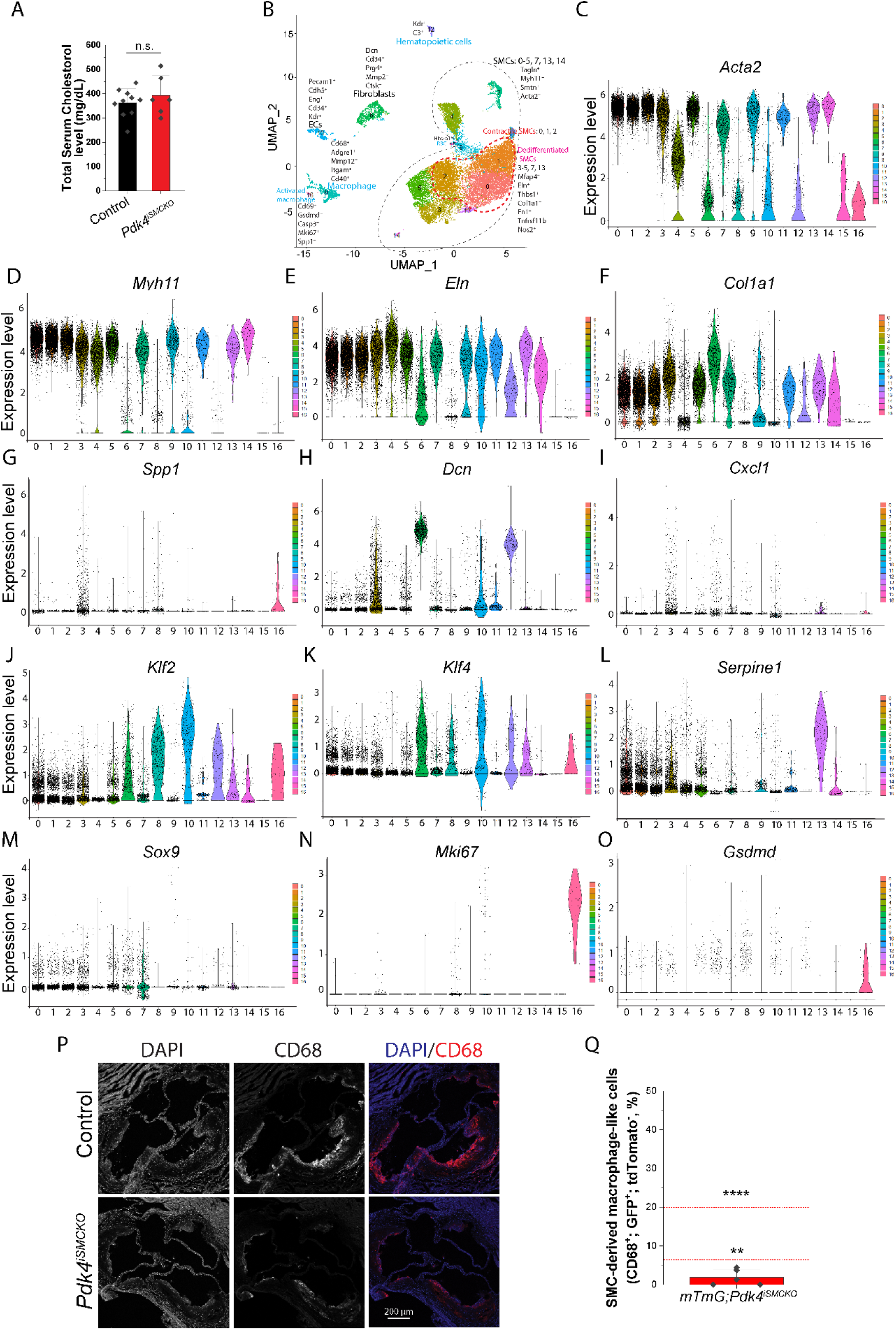
***Pdk4* knockout ameliorates atherosclerosis and maintains SMC contractile phenotype in *ApoE^−/−^* mice.** (A) Comparison of total serum cholesterol levels between control and Pdk4iSMCKO mice after 4-month HFD feeding (**B**) FeaturePlot analyses of aortic cells from both control and *Pdk4^iSMCKO^* mice, with enriched mRNAs in each cluster. (**C-O**) Violin plots showing expressions of key cell differentiation markers among all the clusters. (**P**) Representative images of CD68 staining of aortic root cross-sections from control and *Pdk4^iSMCKO^* mice. (**Q**) Quantification of SMC-derived fraction of CD68^+^ macrophages from five mTmG;*Pdk4^iSMCKO^* mice, with green fluorescence expressing (GFP^+^) and without TdTomato. One sample t-test was performed comparing 5 mice with the reported mean 20% SMC-derived macrophages by Owsiany *et al*. (*32*), or 6.3% as reported by Chen *et al.* (*9*).

### Supplementary Table

**Supplementary Table 1:** Gene expression profile of clusters from scRNAseq.

| Cluster# | 0 | 1 | 2 | 3 | 4 | 5 | 6 | 7 | 8 | 9 | 10 | 11 | 12 | 13 | 14 | 15 | 16 |
| --- | --- | --- | --- | --- | --- | --- | --- | --- | --- | --- | --- | --- | --- | --- | --- | --- | --- |
| SMC/Contractile | +++ | +++ | +++ | +++ | ++ | +++ | - | ++ | - | ++ | - | ++ | - | ++ | ++ | - | - |
| SMC/Synthetic | + | + | + | +++ | + | ++ | +++ | ++ | - | + | - | ++ | - | ++ | + | - | - |
| Macrophages | - | - | - | - | - | - | - | - | +++ | - | - | - | - | - | - | - | +++ |
| Innate Immune | - | - | - | ++ | - | - | ++ | - | + | - | - | - | - | + | - | - | + |
| Fibroblasts | - | - | - | + | - | - | +++ | - | - | - | + | + | ++ | - | - | - | - |
| Endothelial cells | - | - | - | ++ | - | - | + | + | - | - | +++ | - | - | + | - | - | - |
| Osteoblast-like | - | - | - | ++ | - | - | + | + | + | - | - | - | - | - | - | - | + |
| Stem cell like | - | - | - | + | - | - | ++ | - | + | - | + | - | +++ | - | - | - | + |

+++ strongest expression, ++ medium, + low, - No expression
Contractile SMC clusters: 0, 1 and 2
Synthetic SMC clusters: 3, 4, 5, 7, 9, 11, 13 and 14
Macrophage clusters: 8 and 16
Fibroblasts: 6
Endothelial cells: 10
Hematopoietic cells: 12
Red blood cells: 15

**Supplementary Table 2:**
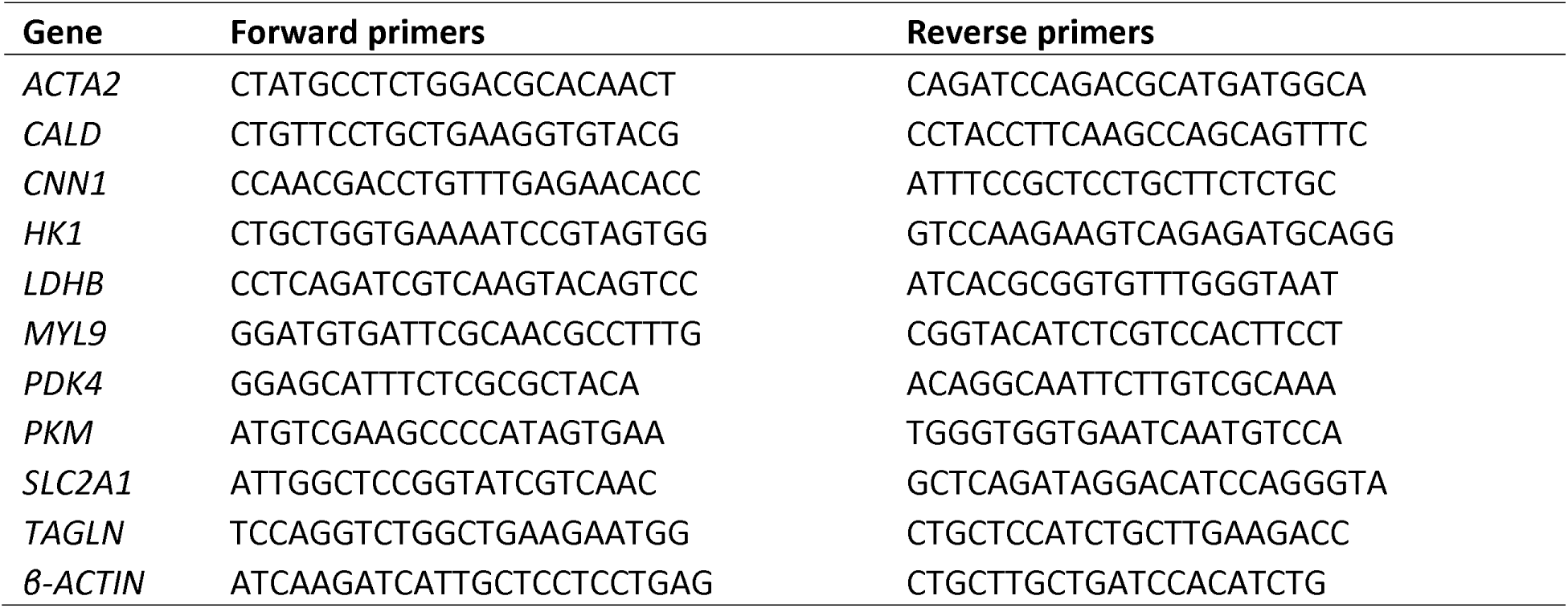
Sequences (5’-3’) of primers used in qPCR.

